# Functional profiling of spacecraft cleanroom microbiomes through genome-wide phenotype predictions

**DOI:** 10.64898/2026.08.28.747777

**Authors:** Alexander Mahnert, Tobias Medicus, Christina Kumpitsch, Christine Moissl-Eichinger, Jonathan Carter, Mark A. Sephton, Silvio Sinibaldi, Petra Rettberg

**Affiliations:** Medical University of Graz, Diagnostic & Research Institute of Hygiene, Microbiology and Environmental Medicine, Graz, Austria; University of Vienna, Computational Systems Biology, Vienna, Austria; Coventry University, Centre for Fluids and Complex Systems, Coventry, UK; Department of Earth Science and Engineering, Imperial College London, London, UK; ESA-ESTEC, Noordwijk, Netherlands; German Aerospace Center, Institute of Aerospace Medicine, Cologne, Germany

**Keywords:** Planetary Protection, JUICE mission, icy worlds, icy moons, phenotype prediction, functional microbiome, survivability, extraterrestrial environments, machine learning, metagenome assembled genomes

## Abstract

Current planetary protection approaches rely heavily on spore-based tests developed for Mars missions and may not adequately assess contamination risks for icy ocean worlds such as Europa. We developed a genome-based framework combining deep shotgun metagenomics and supervised machine learning to predict survival-relevant microbial traits in ESA JUICE launch-site cleanrooms. From 183 genome bins, 25 representative genomes were analyzed for traits including cryotolerance, desiccation tolerance, salt resilience, anaerobic metabolism, autotrophy, and sporulation. Several skin-associated microbes carried multiple relevant traits, and some appeared actively replicating. A broader meta-analysis of 1,868 genomes showed that trait profiles vary strongly within taxa, demonstrating that taxonomy alone is insufficient for risk assessment. This framework complements current planetary protection assays, helps to predict how microbes would survive in a new biotope, and supports functional, risk-informed contamination monitoring for future space missions.

## Introduction

Preventing forward contamination of extraterrestrial environments is a central principle of planetary protection ^1–3^. To comply with international requirements, spacecraft assembly cleanrooms and hardware are routinely monitored for microbial burden and biodiversity ^4–6^, since cleanrooms harbor low-biomass but persistent taxa, including stress-tolerant lineages relevant to planetary protection ^7–10^. For decades, ESA and NASA have relied on a colony-count assay including a heat-shock as the standard method (ECSS-Q-ST-70-55). This assay serves as an easy to measure proxy for the overall bioburden. In this assay, samples are exposed to 80 °C for 15 minutes, plated on nutrient-rich agar, and incubated aerobically at 32 °C for 72 hours ^11^. While effective in enumerating spore-forming organisms, this approach provides only a limited view of the total spacecraft-associated microbiota. Sporulation is indeed a powerful survival strategy, but many extremophiles potentially relevant for planetary protection do not depend on spores for persistence under harsh space-like conditions ^12,13^.

Key traits for microbial survival and replication in extraterrestrial environments, particularly on icy worlds, extend beyond sporulation ^14^. These include desiccation resistance ^15,16^, tolerance to a wide range of UV and ionizing radiation ^17,18^, oligotrophy ^19^, psychrophily or cryophily ^20–24^, anaerobic metabolism, and halotolerance or halophily. Collectively, these traits could enable organisms to endure desiccation, radiation, low-nutrient availability, subzero temperatures, anoxic conditions, and saline subsurface oceans characteristic of icy moons ^25,26^. Accurate recognition and modeling of these survival strategies are essential for refining contamination risk assessments.

To advance beyond cultivation-based assays, ESA has been investing in the development of tailored molecular approaches for cleanroom monitoring. Since 2003, successive projects (MiDiv 2003–2005, BioDiv 2006–2010) and currently 36 planetary protection campaigns (PP-VERI 2011–2026) have combined cultivation-based spore counts and alternative growth assays with molecular methods. In particular, next-generation sequencing (NGS) of 16S rRNA gene amplicons was implemented from 2016 onwards. This cultivation-independent method offers high-throughput, low-cost insights into microbial community structure, even in low-biomass environments. However, its application in spacecraft cleanrooms also exposes critical challenges, as a current meta-analysis of 205 samples showed extensive overlap between biological samples and controls, with common human commensals appearing in both and no significant differences in alpha or beta diversity despite controls comprising over half the dataset ^27^. Machine-learning models failed to distinguish sample types, and filtering suggested that nearly 7% of reads were contaminants ^28,29^. Obviously, low-biomass environments such as spacecraft assembly cleanrooms are particularly susceptible to background contamination from DNA extraction kits, reagents, and various molecular assays that reach their limit of detection. Hence, it is essential to incorporate appropriate sample controls (field blanks, sampling material, DNA extraction, lab reagents, negative and positive controls, library preparation controls, etc.) throughout all stages of the entire workflow, allowing subsequent bioinformatic statistical corrections based on abundance and prevalence. Clear site-specific patterns also emerged, such as distinct, climate-driven community profiles at ESA’s Kourou facility. These findings demonstrate both the potential and limitations of amplicon sequencing for planetary protection: the method is highly sensitive but constrained by primer bias, missing potential key taxa and changing actual abundances, low phylogenetic resolution ^30^, limited functional insight, and strong susceptibility to contamination.

Given these limitations, whole-genome shotgun metagenomics is emerging as a promising alternative ^31^ as it extends beyond marker-gene community profiling and resolves genome-level functional capacities. Unlike targeted amplicon approaches, metagenomics sequences all available DNA in a sample, reducing bias and enabling simultaneous taxonomic and functional profiling. It offers multi-level resolution, from reads to contigs to metagenome-assembled genomes that could dramatically expand representation of uncultured taxa, relevant when cleanrooms contain atypical, stress-selected organisms that may be hard to culture. Genome-centric metagenomics has the ability to resolve strain-level diversity, single-nucleotide polymorphisms (SNPs), and gene synteny. This genome-centric view enables advanced analyses such as metabolic modeling and phenotype prediction. In addition, genome-centric metagenomics requires sufficient coverage to reconstruct a genome from the metagenome. Consequently, this method is less susceptible to background noise contamination from isolated and scattered sequencing reads compared to simple profiling that tends to produce false positives. However, these advantages come with higher input DNA requirements, increased costs, longer analysis times, computationally demanding workflows, and dependence on representative reference databases. Thus, there are still some challenges to overcome before these methods can actually be implemented in spaceflight^32^.

### Objectives

The overarching objective of this work was to explore how metagenomic and computational approaches can predict microbial behavior in a new biotope and enhance planetary protection monitoring. Specifically, we focused on the prediction of microbial phenotypic traits relevant to survival and replication on icy worlds, using machine learning approaches applied to metagenomic data. By bridging molecular microbiology with advanced computational methods, this study seeks to refine contamination risk assessments and develop a framework that extends beyond taxonomic inventories to understand the functional and phenotypic potential of spacecraft-associated microbiota that specifically complies with future planetary protection guidelines and can reduce the risk of costly false negatives further. (Fig. 1).

**Figure 1:**
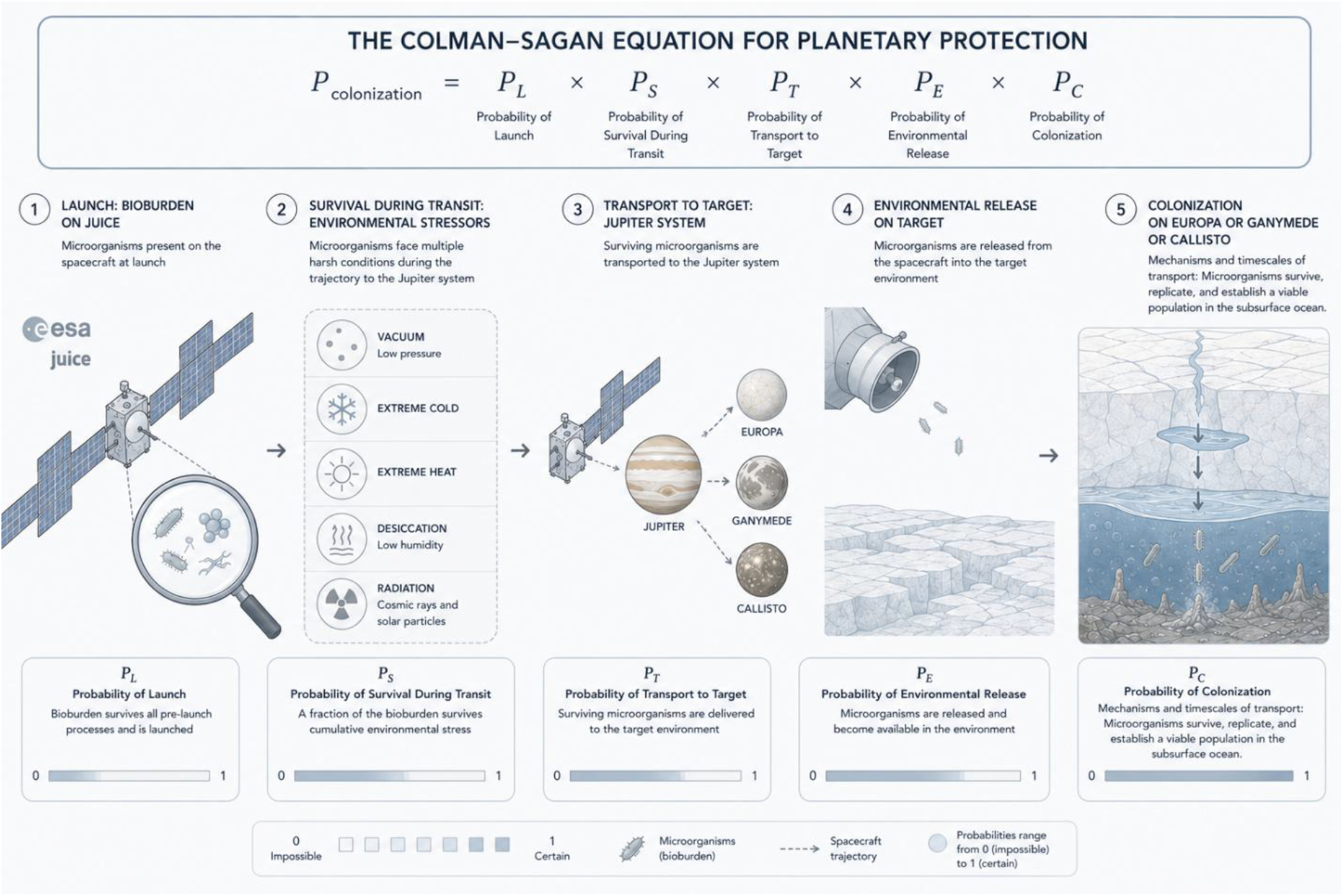
Graphical abstract covering the rationale behind the ProConTra (Probability of Contamination and Transport Mechanisms) project and its use-case of ESA’s JUICE mission to the Jupiter system and how genome-centric metagenomics could improve estimates for the Colman-Sagan Equation for planetary protection. Each step in the equation contributes to the overall probability of colonizing an extraterrestrial target referred to as “forward contamination”. This study focuses on step 1 and 2. Genome-centric metagenomics are used to determine the bioburden at launch (step 1) and the newly developed genome-scale phenotype prediction tools can assess the survivability in face of diverse environmental stressors (step 2). The probabilities for transport and release of harmful entities during the trajectory are addressed in step 3 and 4 as presented in ^33^. Finally, step 5 shows that the colonization of an icy ocean world depends on respective transport mechanisms and timescales from the surface to the subsurface ocean as well as the capabilities of microbes to survive, replicate and establish a viable population in the new environment.

## Methods

### Sampling

All available floor surfaces of cleanrooms S5C and S5A in Kourou were sampled to retrieve enough biomass for shotgun metagenomics. The entire floor surface of each cleanroom was quartered to gain replicates of each room for proper statistics and improved co-binning of metagenome-assembled genomes (MAGs). Throughout the process, DNA-free gloves (abf diagnostics, Germany) were worn, and DNA-free equipment was used. The equipment was either decontaminated by sodium hypochlorite solution (0.6%) for 15 minutes and UV (254 nm) sterilization or baked at 220°C for 24 h. For the sampling procedure itself, the floor squeegee was placed flat on the surface and rubbed over the entire surface using a firm, steady pressure. The same sample area was rubbed with the wipe a total of three times, rotating the direction of motion first 90 degrees and then 135 degrees. If necessary, wipes were remoistened with PCR-grade water that was sprayed directly on the surface with an atomizer. In this sampling campaign, surfaces of approx. 87.25 m^2^ in S5C and 75 m^2^ in S5A were sampled. Lastly, the wipe was placed into a sterile transport tube. The wipes were stored in a Styrofoam box with cooling packs at +4°C till transport to the laboratory at the Medical University of Graz, and then stored at −20°C until further processing. For each room, a field blank (negative control) was taken. The field blanks were collected by removing a sterile sampling wipe from its tube, opening the wipe and touching it with gloves, and placing it back into the tube. Back at the laboratory, also extraction blanks were used to control for molecular contaminations of reagents and equipment. All controls were processed and analyzed in the same way as biological samples. Additionally, negative and positive controls were included during sequencing library preparation and shotgun metagenomics sequencing.

### DNA extraction

For extracting microbial cells, wipes were taken out of the freezer (−20°C) and thawed overnight in the fridge (+4°C). Thawed wipes were placed using heat sterilized (at +220°C for 24 h) tweezers in sterilized 250ml glass bottles, and 100ml of PCR-grade water (LiChrosolv®, Supelco) was added to each bottle. The bottles were then placed in an ultrasonic bath so that the water level in the bath was higher than in the bottles. The bottles were sonicated for 2 x 5 minutes and vortexed at maximum speed for 1 minute. The liquid was concentrated to 200 µl using UV (254 nm) sterilized Amicon filters (Amicon Ultra-15 Centrifugal Filter Units, Ultracel 50K, Merck, Germany). The UV (254 nm) sterilization was performed in a clean bench by opening the Amicon filters and exposing them to UV (254 nm) radiation for 1 hour. Concentrating the liquid from 100 ml to 200 µl was accomplished by serial centrifugations of 10 ml aliquots at + 4 °C and 4000x g for 10 minutes. DNA was extracted from the remaining liquid using the FastDNA Spin kit (with Lysing Matrix E-tubes; MP Biomedicals, Germany) according to the manufacturer’s instructions. The amount of DNA was measured with the Qubit HS assay (Thermo Fisher Scientific, Austria). DNA concentration was below the detection limit in all samples.

### Library preparation and shotgun sequencing

Shotgun sequencing libraries of all samples using the NEBNext® Ultra™ II FS DNA Library Prep with Beads (24rxn) were prepared by the Core Facility for Molecular Biology at the Medical University of Graz, including tests for the library preparation to optimize the fragmentation time and to check that samples will deliver successful libraries before their shipment to the sequencing facility VBCF (Vienna BioCenter Core Facility). At the VBCF, the library was sequenced on a 2x NovaSeq SP500 cycles Flowcell with an intended output of 1.3 – 1.6 billion reads (1.92 – 2.34*10^8^ reads per sample and Flowcell).

### Data analyses

On average, 2.24*10^8^ raw shotgun sequences were produced per sample. All reads were analyzed in a gene- and genome-centric approach. To allow best bioinformatic reproducibility we used metagenome-atlas 2.18.0 ^34^, a Snakemake workflow that concatenates all necessary steps from quality control (removal of PCR duplicates, quality trimming of reads, removal of common contaminants like adapter sequences, phiX and human host reads), metagenome assembly (error correction, merging of paired-end reads, assembly with metaSpades and post-filtering), genomic binning with Metabat2 ^35^, assessment of genome quality with CheckM2 ^36^ and GUNC ^37^, refining bins with DASTool ^38^, dereplication of bins with dRep ^39^ and Skani ^40^, taxonomic classification according to GTDB ^41,42^, strain resolved genomics achieved with inStrain analysis ^43^, quantification of genomes, and functional annotation of genomes and contigs with eggNOG ^44,45^ and DRAM ^46^, and clustering of redundant genes with linclust ^47^. The following tools were used for this workflow: bbmap: 39.06, bowtie2: 2.5.1.3, checkm2: 1.0.1, das_tool: 1.1.6, diamond: 2.1.9, dram: 1.4.6, drep: 3.5.0, eggnog-mapper: 2.1.12, fastani: 1.34, fraggenescan: 1.31, gtdbtk: 2.3.2, hmmer: 3.4, inStrain: 1.5.7, mash: 2.3, maxbin2: 2.2.7, megahit: 1.2.9, metabat v2, minimap2: 2.26, mmseqs2: 15.6f452, prodigal: 2.6.3. In addition, for rapid gene-centric taxonomic profiling, we used Kraken2 ^48^ (confidence cut-off 0.3) and quantification of species with bracken (min. number of reads = 100) according to the GTDB and PlusPFP database to cover Eukaryotes and viral information besides Prokaryotes.

### Controls

All controls were processed and sequenced alongside study samples to enable contamination detection, performance verification, and reproducible interpretation. Controls were carried through the complete workflow (sampling, extraction, library preparation, sequencing) and analyzed in parallel during bioinformatic processing. Field/sampling blanks (sterile wipes, filters, and transport containers exposed to the sampling environment but without intentional collection and contact to the surface of interest), extraction blanks (molecular-grade water or buffer processed through the full extraction protocol), and no-template controls (NTCs; library reactions lacking sample DNA) were included to monitor environmental, kit-derived, and reagent contamination, respectively. Positive controls (known genomic DNA) were included to verify assay performance, monitor extraction and library-preparation efficiency, and permit quantitative assessment.

### Machine learning and prediction of PP-relevant phenotypes

Three new classifiers were developed in the frame of this project to predict halo resilience, cold resistance, and desiccation resistance. The halo resilience classifier was trained using data from both halophilic and halotolerant organisms and hypersaline habitat-associated organisms, leading to the model being termed “haloresilient” (Table 1). In the process of developing our machine learning models, careful consideration was given to rules for the selection and refinement of the dataset. The idea was to establish parameters with a stark contrast between the positive and negative sets, based on environmental conditions and growth characteristics. As shown in Supplementary Table 1, these encompass, for example, the haloresilient model selected species with the ability to grow above 10% NaCl for the positive set and growth only below 3% for the negative set. The decision was made to create a grand enough distinction between the positive and negative sets to enable the algorithm to potentially more accurately detect and distinguish the specific traits, especially the extremes. These parameters were used for the fine-tuning and discrimination of data during the refinement step; see Supplementary results in Supplementary Information. Furthermore, for the desiccation resistance classifier, the habitual constraint of whether the organism was known to exist in desert environments or not is termed the “desert trait predictor,” emphasizing adaptations specific to desert conditions. However, we are aware that not all organisms in desert environments are desiccation resistant and that not all desiccation resistant organisms will be able to survive for several years in space vacuum during the journey. Further details on the methodology behind developing and refining the machine learning models, how to evaluate them, the formulas used, and simulations of prediction accuracy in dependence on genome completeness and contamination can be found in the Supplementary Information. Phenotype predictions of all available trait models were conducted with phenotrex v0.6.0 (https://phenotrex.readthedocs.io/en/latest/usage.html, https://github.com/LokiLuciferase/phenDB?tab=readme-ov-file, ^49^ at a minimal probability of 0.5, revealing the top 10 most predictive features for selected traits. Required input files of eggNOG cluster IDs were generated with a custom workflow including gene prediction with prodigal v2.6.3 ^50^, taxonomic classification with GTDBtk v2.4.1 r226 ^41^, and using PyHMMER ^51^ instead of deepNOG ^52^ in a parallelization mode for better time scaling and improved sensitivity and specificity. See the GitHub repository for useful scripts and details (https://github.com/Mechah/ProConTra).

**Table 1:** Model details, including hyperparameters, including balanced accuracy of the final model iterations with cross-validation using 5 folds and 10 replications.

| model | genomes | cross-validated balanced accuracy | n estimators | max depth | gamma | min childweight | colsample by tree | subsample |
| --- | --- | --- | --- | --- | --- | --- | --- | --- |
| Haloresilient2 | 560 | 0.8248 ± 0.0324 | 100 | 4 | 0.5 | 9 | 0.4 | 0.6 |
| Cryotolerance5 | 184 | 0.8006 ± 0.0636 | 80 | 3 | 0.5 | 4 | 0.5 | 0.5 |
| Desiccation2 | 638 | 0.7684 ± 0.0386 | 80 | 6 | 1 | 4 | 0.8 | 0.8 |

### Meta-analyses

Phenotype predictions from the JUICE mission at ESA’s spaceport in Kourou were assessed in a broader context and compared to five selected genome collections (https://www.dsmz.de/collection/catalogue/microorganisms/special-groups-of-organisms/esa-strains) ^53–56^. This meta-analysis covered 1,868 genomes/bins and 78,456 individual predictions. The analyses included a detailed multi-study comparison of planetary protection-relevant phenotypes and a focused analysis of the genus *Staphylococcus*. Statistics and visualizations were conducted in R using the packages: readr, dplyr, ggplot2, rstatix, writexl, purrr, tibble, stringr, ggpubr, tidyr, and forcats.

## Results

### Phenotypic models

#### Cross-Validation

Cross-validation was used to evaluate the performance of our initial phenotypic models. As shown in Table 1, our newly developed models achieved a balanced accuracy of around 0.76 to 0.82. Notably, Desiccation resistant2 emerged here as the least accurate out of the three models, with the lowest Balanced Accuracy. On the other hand, Cryotolerant5 seems to be the least robust with the highest variance.

#### Random Split Evaluation

The performance of our models was further investigated by random splits of the dataset. The random split method divides the dataset into training and testing sets randomly, typically in an 80/20 ratio, ensuring that the models are evaluated on unseen data. The histogram in Supplementary Fig. 1 and Table 2 captures the accuracy of all models at confidence thresholds of 0.5 and 0.7.

**Table 2:**
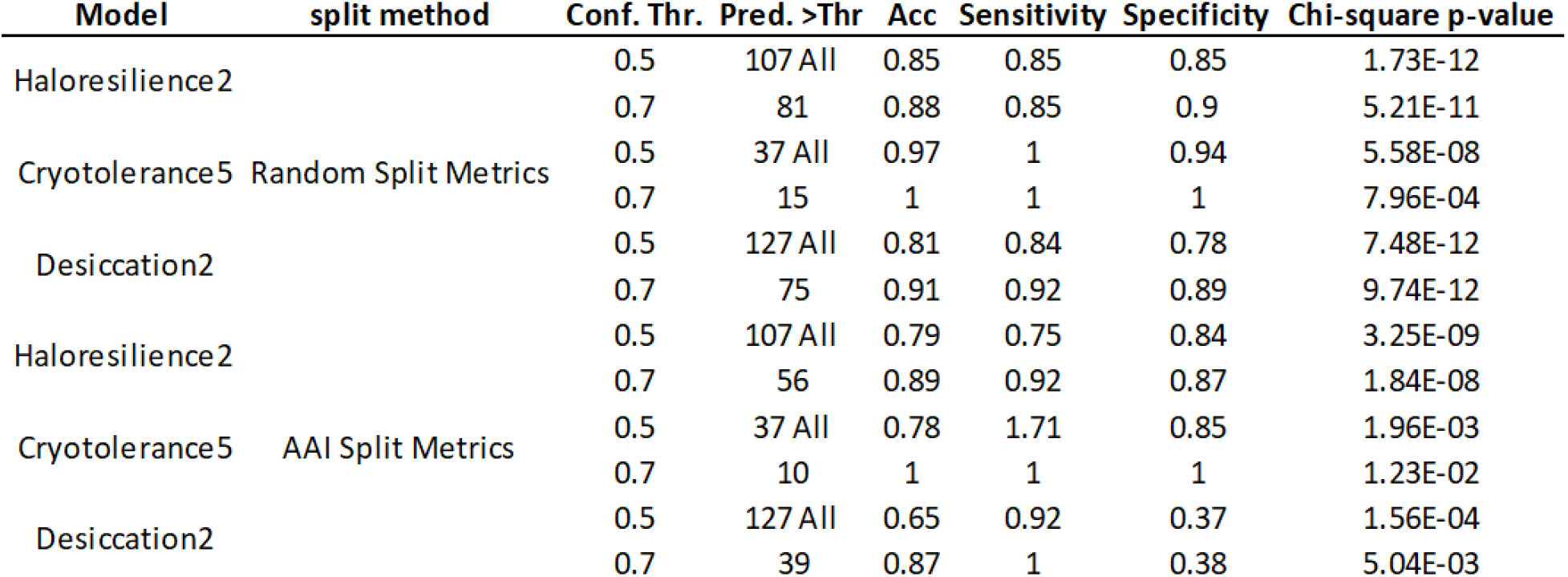
Model performance at different split methods and confidence thresholds. Performance metrics of Haloresilience2, Cryotolerance5, and Desiccation2 models at confidence thresholds (Conf. Thr.) of 0.5 and 0.7. ‘Pred. *>*Thr’ denotes the number of predictions above the threshold, with ‘All’, which means all predictions are above 0.5. Accuracy (Acc), Sensitivity, and Specificity are provided for each threshold. The Chi-square p-value evaluates the prediction accuracy against random guessing, confirming the statistical robustness of the models.

Supplementary Fig. 1 and Table 2 show a notable increase in classification accuracy when the confidence threshold is raised from 0.5 to 0.7. This all comes at the price of disregarding predictions, considerable amounts, ∼24% for Haloresilience2, ∼59% for Cryotolerance5, and ∼41% for Desiccation2, of the predictions are below 0.7. Interestingly, the accuracies of our models fall within the balanced accuracy interval presented in Table 2, except for the cryotolerance model. Further evaluation is presented and described in the Supplementary Information. This evaluation is based on an AAI split to better cover the diversity of input genomes. It also includes estimates of the effects of genome completeness and contamination levels, as well as details of the overall refinement process.

### Confidence of Models

The confidence levels across the various classifiers were analyzed via the random split method and were visually summarized in Supplementary Fig. 2. This graphical representation provides an intuitive comparison of the model’s confidence in its predictions. When examining the box plot, we observe that the sporulation classifier, derived from data processed through PhenDB (Supplementary Table S16) ^49^ and adapted to our workflow, exhibits a remarkably high median confidence score and density. Its outlier points, though numerous, still maintain a level of confidence that does not exceed the upper quartile of the other models, indicating strong predictive certainty. The Haloresilient2 classifier displays the best range and highest median in confidence scores in comparison to the other newly created models. Despite the breadth of Haloresilient2’s confidence interval, no model approaches the confidence level demonstrated by the Sporulation classifier. This stark contrast in confidence levels could be attributed to the specificity of traits associated with sporulation, which may be more distinct and easier for the model to identify compared to the broader and more varied traits associated with cryotolerance. Our specifically trained and evaluated phenotype prediction models for icy worlds were then applied in a use case on metagenomic data obtained from the JUICE mission sampling campaign.

### JUICE Kourou data description

Quality filtering of metagenomic data obtained from the sampling campaign in Kourou retained on average 84.3±8.2% of all reads per sample. Still, a high proportion of reads (63±14.7% on average) could be aligned to the human genome and were removed from downstream processing. *De novo* assembly with metaSpades resulted in contigs with an average N50 of 6.17*10^3^±3.75*10^3^. Binning these contigs into draft genome bins with Metabat2 allowed the recovery of 183 bins (Fig. 2A). Scoring, refining and dereplication of bins resulted in 25 representative MAGs of sufficient quality (average completeness: 93±8%, contamination: 1±1%, genome size: 2.86*10^6^±1.97*10^6^, N50: 8.16*10^4^±8.8*10^4^, number of contigs: 494±640, coding density: 0.9±0.04, gene length: 308±22, GC content: 05±0.1). MAGs were then annotated as described in our Method section. It is also important to mention that the obtained genomes only represent 18.1±17.6% (minimum 6.6%, maximum 57.4%) of the complete data set, and thus only a minor fraction of the microbial diversity of sampled cleanrooms in Kourou (Fig. 2B). Hence, gene-centric taxonomic profiling approaches (e.g. with kraken2/bracken against the PlusPFP database) were processed in parallel to shed also light on the microbial diversity not represented by a MAG. This analysis revealed that most signatures resulted from common skin commensals like *Cutibacterium acnes*, *Corynebacterium*, *Staphylococcus,* or *Micrococcus luteus*, fungi like *Malassezia*, and plants like *Musa acuminata*, *Lactuca sativa,* or *Triticum aestivum* (Supplementary Fig. 9). Focusing on the obtained genomes, the top 10 most abundant genomes in our dataset were *Cutibacterium acnes* (3.64*10^4^), *Iningainema* (1.37*10^4^), *Romboutsia*_A (8.98*10^3^), *Lawsonella clevelandensis*_A (8.77*10^3^), *Lentilactobacillus senioris* (4.24*10^3^), *Aliterella* (4.78*10^3^), *Chrocococcidiopsidaceae* (5.09*10^3^), *Corynebacterium tuberculostearicum*_C (3.2*10^3^), *Leptolyngbyaceae* (3.05*10^3^), and *Haematomicrobium sanguinis* (2.27*10^3^) (Fig. 2C and 2D). Comparing our data on the sub-species level using inStrain, we identified *Cutibacterium acnes* (min. ANI 100, min shared genome coverage 99.99%), *Staphylococcus epidermidis* (min. ANI 99.993, min shared genome coverage 93%), and *Cutibacterium namnetense* (min. ANI 99.92, min shared genome coverage 93%), as not only having high similarity, but also having higher genome coverage among taken samples than *Haematomicrobium sanguinis* (min. ANI 99.996, min shared genome coverage 65%), *Staphylococcus warneri* (min. ANI 99.9, min shared genome coverage 60%), *Corynebacterium tuberculostearicum*_C (min. ANI 99.986, min shared genome coverage 80%). This could indicate greater subspecies diversification between the two sampled cleanrooms for the latter representative genomes.

**Figure 2:**
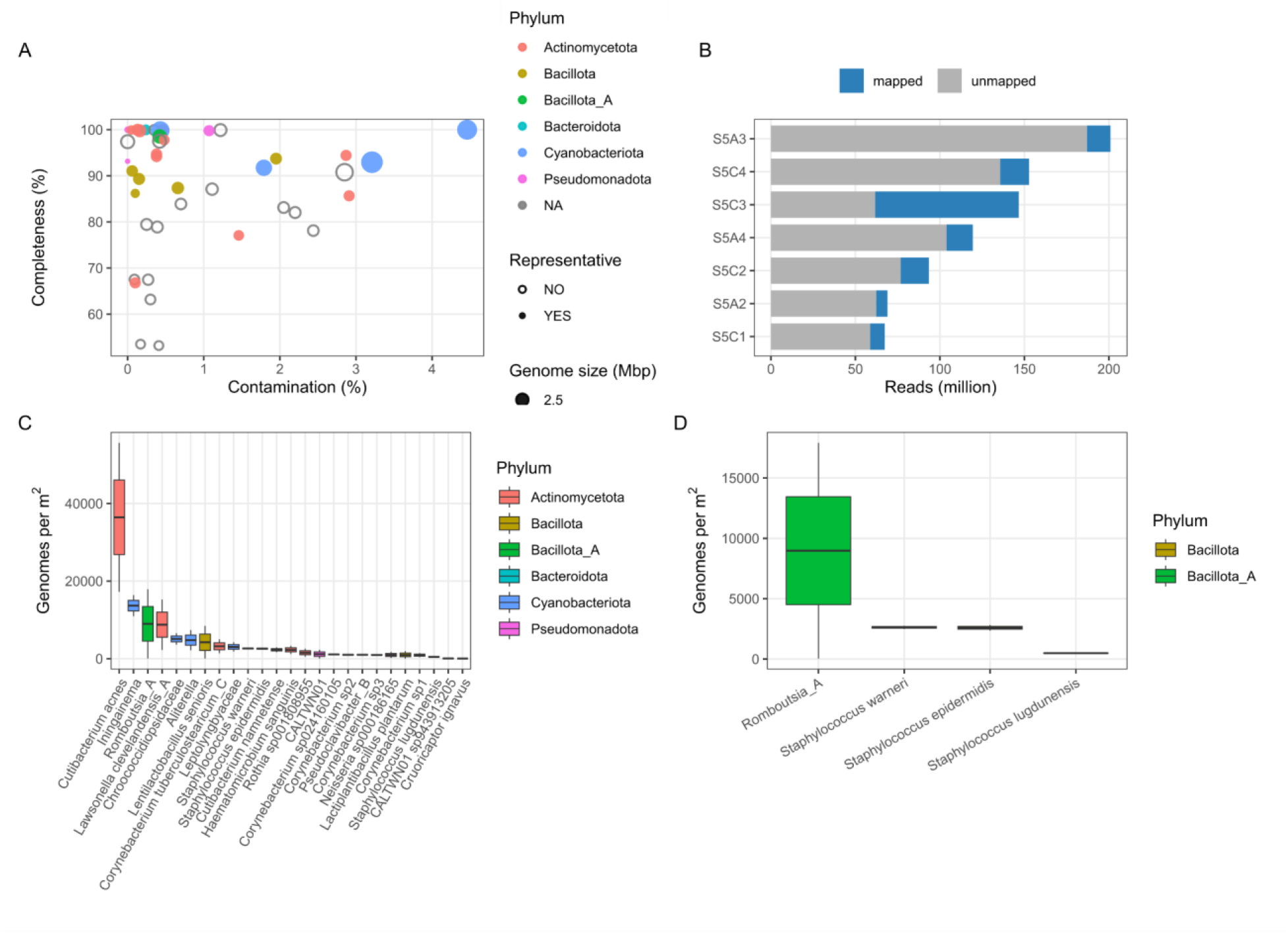
JUICE use-case Kourou genome-centric data description **A**) Scatterplot of bin quality estimates (completeness and contamination), major phyla, and representative metagenome assembled genomes (MAGs). **B**) Proportion of data representation by MAGs displayed as a stacked bar chart of MAGs mapped on raw QC reads. **C**) Boxplot of most abundant MAGs per m^2^. **D**) Abundance of taxa showing planetary protection-relevant phenotypic traits.

### Phenotype predictions

A major interest of our research was the prediction of PP-relevant phenotypic traits (see Materials & Methods for more details). From our list of 25 representative MAGs, 19 showed an anaerobic trait, 6 MAGs showed an autotrophic lifestyle, 2 MAGs were cryo-tolerant, 8 MAGs were desiccation resistant, 5 MAGs were halo-resilient, and 1 MAG was predicted to have the capability to form spores. According to the phenotypic trait prediction, *Staphylococcus warneri, S. lugdunensis, S. epidermidis*, and *Romboutsia*_A showed three or more predicted traits that were assessed as relevant for Planetary Protection and therefore could be considered to have the highest probability of surviving the journey to Jupiter’s icy moons (Fig. 3 and Fig. 4). However, it is important to note that only a combination of resistance mechanisms against all stress conditions should be considered as being critical for survival. For example, if a microorganism is not resistant to desiccation, all other traits would consequently be then irrelevant for PP. *S. warneri, S. lugdunensis*, and *S. epidermidis* were predicted to be anaerobic, desiccation-tolerant, halo-resilient, having a fermentative and saccharolytic lifestyle, are Gram-positive, and produce acetoin and lactic acids; while *S. epidermidis* showed additional predicted traits for cryotolerance and nitrogen fixation. *Romboutsia*_A was predicted to be an anaerobe, cryotolerant, having the capability to form spores with a fermentative and saccharolytic lifestyle, fixing nitrogen, is Gram-positive and motile, and produces acetoin, butyric acid, formic acid, and hydrogen. Most of these predicted traits are in agreement with the current literature. Nevertheless, for instance the predicted acetoin production for the staphylococci is plausible but may be strain-dependent. Cryotolerance is not a standard trait for staphylococci and could show adaptation to the cleanroom environment. Nitrogen fixation is not supported by the established literature and could be validated by respective *nif* gene clusters. Predictions of motility, cryotolerance, nitrogen fixation, and specific products such as butyrate, formate, acetoin, and hydrogen should be checked for *Romboutsia*_A and might require experimental validations. It is important to emphasize that, while these species showed positive predictions for many relevant traits, they were not among the most abundant ones. *Rombutsia*_A was more abundant in S5A than S5C (1.79*10^4^ vs. 3.64*10^1^ genomes per m^2^), while the staphylococci were all in a similar range of ∼1.9*10^3^ genomes per m^2^ (Fig. 2C and 2D). As the number of chromosome/genome copies relates to an organism’s growth stage (e.g. 1-4 for staphylococci), we also examined individual growth rates of the representative genomes in an additional analysis. To estimate the potential activity (viability) of the representative MAGs we predicted growth rates from peak to trough estimates (PTR) with GRiD (Fig. 3B) and growth conditions with GenomeSPOT (Fig. 3C). These calculations indicated that 16 MAGs had GRiD values above 1.02 and could be considered being actively replicating at the time of sampling. *Corynebacterium*, *Neisseria*, and *Lentilactobacillus senioris* had even a GRiD value above 2, while the staphylococci were only in a range of 0 – 1.38 GRiD values. However, based on these analyses, it is not possible to determine whether higher replication rates are associated solely with the time at which microbial cells are deposited from human skin onto cleanroom surfaces. Predicted optimal growth conditions from GenomeSPOT were, on average in a range of pH 7±1, salinity 3±2%, and temperature 30±5 °C, and most MAGs were predicted to be tolerant to oxygen (92%). Focusing on the extremes, *Haematomicrobium sanguinis* was predicted to grow in alkaline conditions (pH 10), *Lawsonella clevelandensis*_A at an acidic pH of 4, *Staphylococcus warneri* growing in highly saline conditions (13%), while several species did not require salt for growth, *Corynebacterium* sp024160105 might grow up to 45 °C, and a MAG classified as Alphaproteobacteria CALTWN01 was predicted to grow at only 9°C.

**Figure 3:**
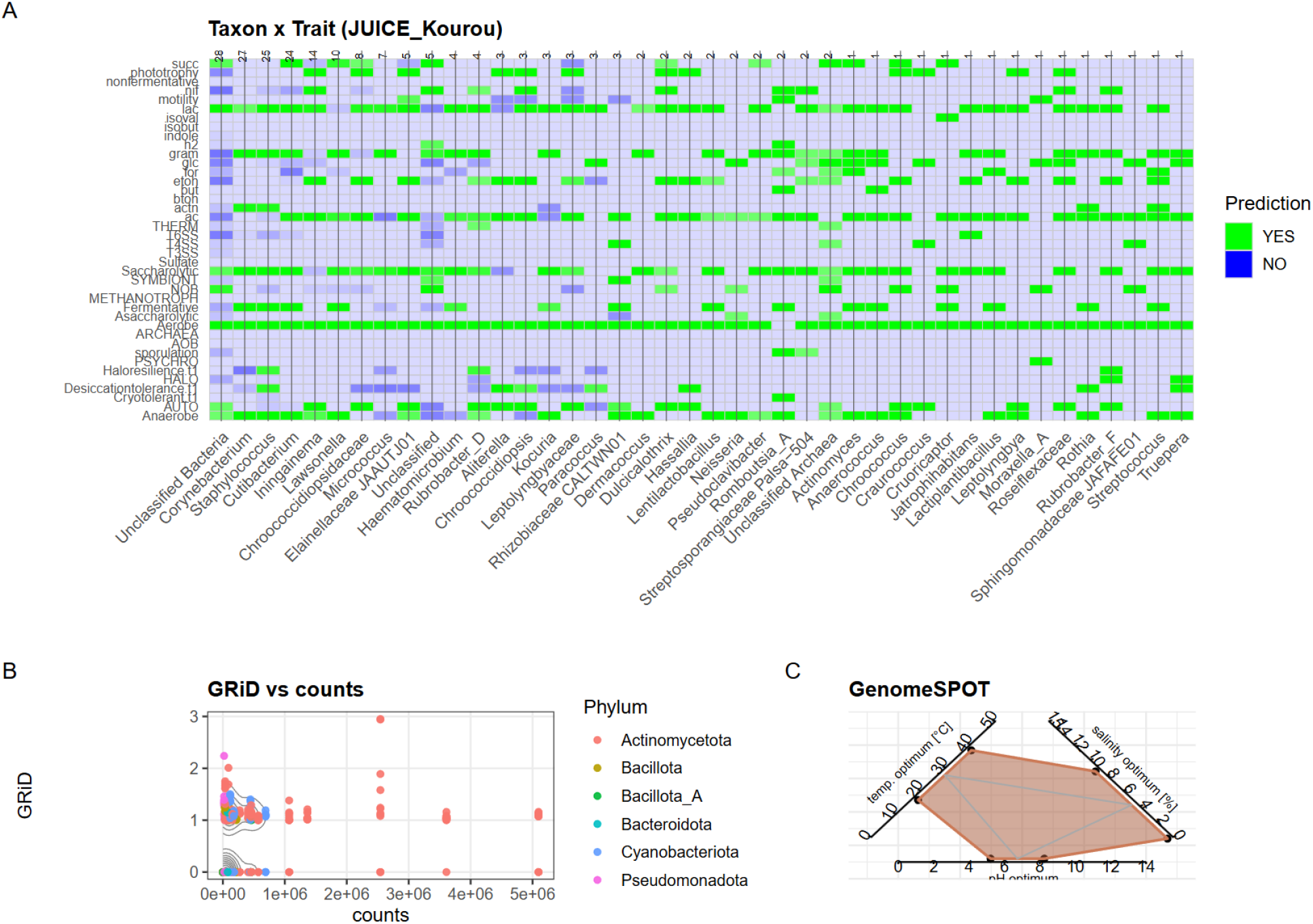
Predicted traits. **A**) Matrix plot showing all predicted phenotypes of all taxa from JUICE Kourou. **B**) Contour plot of predicted growth rates and counts colored according to phylum level. **C**) Polygon plot of predicted optimal growth conditions of all MAGs.

**Figure 4:**
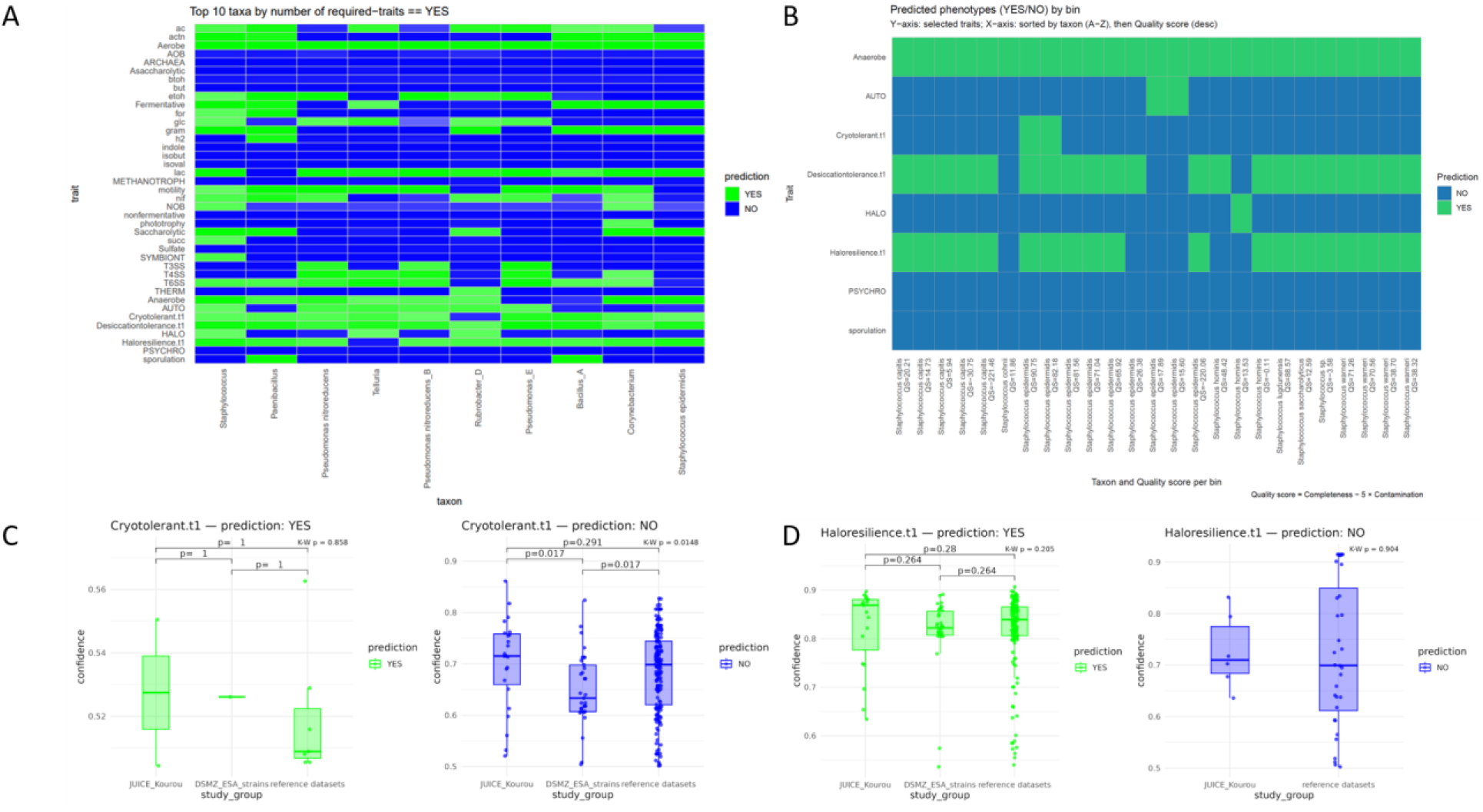
**A**) Matrix plot showing predicted phenotypes for the top 10 most abundant taxa. B) Matrix plot showing only Planetary Protection-relevant phenotypic traits for the most abundant taxon, *Staphylococcus*. **C**) Box plot of confidence scores for exemplary traits.

### Meta-analysis

We then conducted an extensive comparative meta-analysis (see Methods for details) of 1,868 genome bins and isolates leading to 78,456 predictions to place our observations from the JUICE sampling campaign into a larger context (Supplementary Tables 2 - 16). We included five selected genome collections in this meta-analysis: i) Genomes from our ESA strain collection (https://www.dsmz.de/collection/catalogue/microorganisms/special-groups-of-organisms/esa-strains); ii) our ensemble binning approach from various spacecraft assembly cleanrooms ^54^; iii) our spatial and longitudinal genome collection from the International Space Station ^55^; iv) the reference genome collection of the human skin microbiome ^56^; v) and our genome collection of a very deeply sequenced human skin microbiome ^53^. Surprisingly, this vast analysis revealed similar proportions at a high overall median confidence of 0.885 across all PP-relevant phenotypes in other reference datasets (average NO prediction = 79.3%, average YES prediction = 20.7%; see Methods for details). Within this vast meta-analysis, the predictions of PP-relevant phenotypic traits were in a similar range, also for the JUICE mission at the Kourou launch site. The JUICE dataset was characterized by the lowest median confidence score of 0.793. Significant pairwise differences were observed for individual traits, but these are likely a technical tradeoff only resulting from different proportions of low-quality bins, up to complete circularized genomes from isolates covered by other selected reference datasets. Among the PP-relevant phenotypic traits, genomes from the ISS stood out for the anaerobic lifestyle (75.1%, 444/622, chisq.test p = 1.24E-15), the ensemble binning dataset for cryotolerance (25%, 11/44, chisq.test p = 1.28E-6), and ESA’s collection of isolates (https://bacdive.dsmz.de/collection/esa) for desiccation tolerance (76.2%, 99/130, chisq.test p = 1.01E-24), haloresilience (56.9%, 74/130, chisq.test p = 1.72E-16), and sporulation (13.1%, 17/130, chisq.test p = 2.42E-7). These observations correspond very well with the respective characteristics of the selected datasets: According to our analyses, for example, there was a dispersal of fecal-associated microbes from the WHC (Waste and Hygiene Compartment) on the ISS, leading to higher proportions of facultative anaerobes ^55^. Cleanroom environments typically have a lower temperature than various regions of the human body, which are primarily responsible for the contamination of these environments. And the ESA strain collection consists mainly of isolates obtained primarily through the PP standard assay, which relies on the selective enrichment of spore-forming microbes. Focusing on the JUICE dataset and its relative proportions of positive traits of all taxa compared with the other selected studies in our meta-analyses, the JUICE dataset revealed higher proportions of autotrophy (34.4%, 73/212, chisq.test p = 1.06E-14), halophilic (4.7%, 10/212, chisq.test p = 0.177), nitrite-oxidizing bacteria (NOB, 20.8%, 44/212, chisq.test p = 8.77E-19), nif gene clusters (26.4%, 56/212, chisq.test p = 9.9E-19), symbiont lifestyle (4.7%, 10/212, chisq.test p = 0.261), saccharolytic lifestyle (75.5%, 160/212, chisq.test p = 8.88E-10), thermophilic (2.4%, 5/212, chisq.test p = 0.07), phototrophy (25.5%, 54/212, chisq.test p = 3.83E-61), and succinate metabolism (32.5%, 69/212, chisq.test p = 1.8 E-13). Next, we took a closer look at taxon-specific trends: *Staphylococcus, Paenibacillus, Pseudomonas, Telluria,* and *Rubrobacter* showed five or even more positive PP traits, with *Staphylococcus* leading the list of the top 10 taxa. However, among 25 different staphylococci, functional heterogeneity of these PP-relevant phenotypes was evident. These observations underline the need to think about the risk of forward contamination beyond the limits of taxonomic genus or species level boundaries. In comparison to our reference datasets, staphylococci from Kourou during the JUICE mission showed higher proportions of autotrophy (chisq.test p = 0.187), cryotolerance (chisq.test p = 0.574), and a halophilic phenotype (chisq.test p = 0.0128). (Fig. 5).

**Figure 5:**
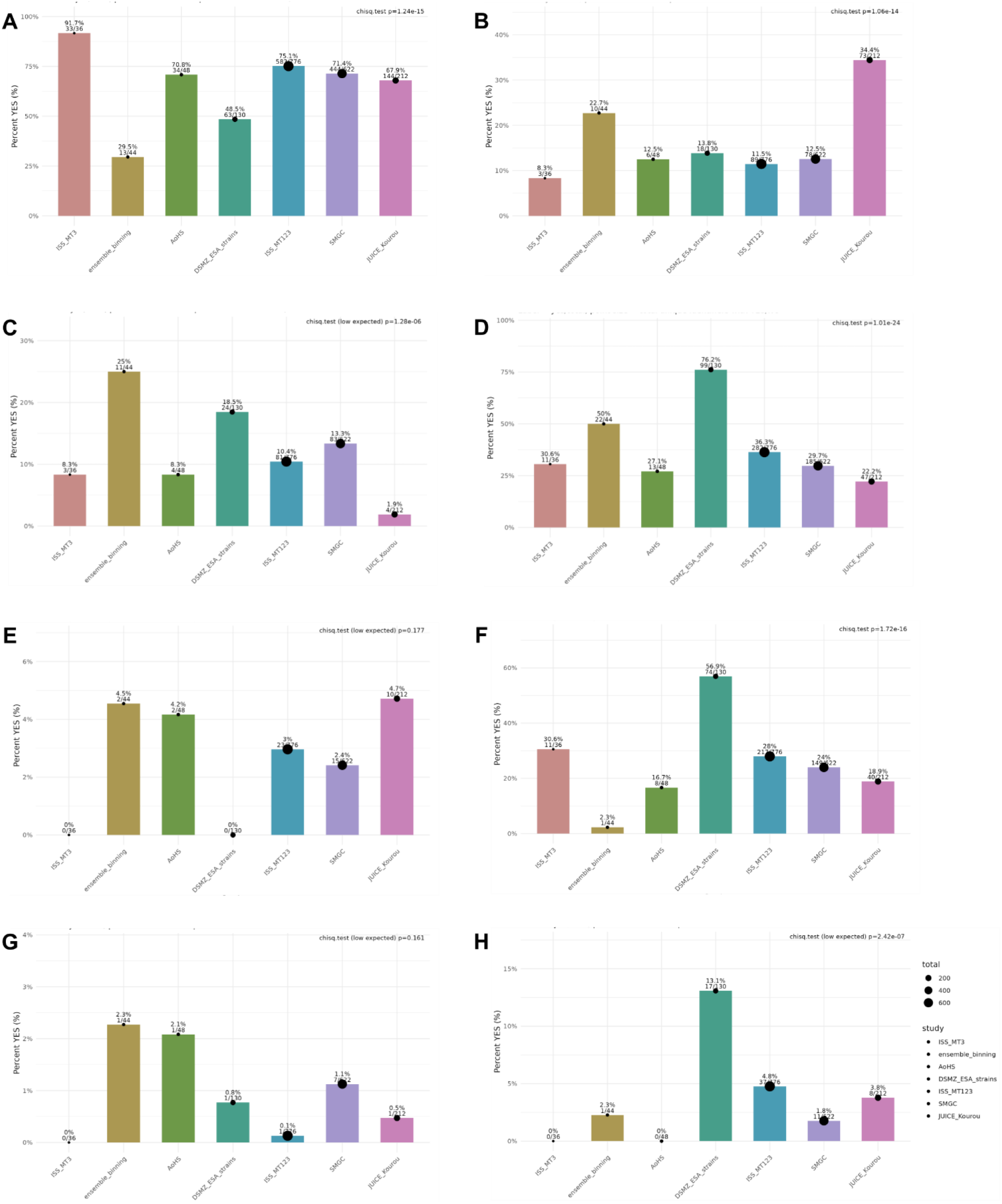
Meta-analysis showing proportion of positive Planetary Protection relevant traits (Y-axis) in comparison to other key references (along X-axis). A) anaerobe, B) autotroph, C) cryo-tolerant, D) desiccation tolerant, E) halophilic, F) halo-resilient, G) psychrophilic, H) sporulation. Included studies: ISS_MT3 (representative MAGs from microbial tracking 3 onboard the International Space Station ^55^); ensemble binning (representative MAGs from cleanroom meta-study ^54^); AoHS (genome bins from deeply sequenced healthy skin microbiome ^53^); DSMZ_ESA_strains (selected genomes of ESA’s strain collection at DSMZ https://www.dsmz.de/); ISS_MT123 (genome bins from all microbial tracking missions 1,2, and 3 onboard the International Space Station ^55^); SMGC (reference collection of genomes and isolates of the skin microbiome ^56^); JUICE_Kourou (genome bins from the JUICE sampling campaign – this study). Proportions including percentage and number of genomes, bins, or isolates are indicated above each bar and also visualized by the increased diameter of a black circle. For each trait the result of a Chi-squared test is displayed in the upper right corner of respective panels.

Finally, for the selected PP relevant traits (anaerobe, autotrophy, cryotolerance, desiccation tolerance, halophilic lifestyle, haloresilience, psychrophilic lifestyle, sporulation), we calculated the most predictive protein features from JUICE Kourou to identify potential targets for future functional screenings (Supplementary Tables 8-15). Prediction of the anaerobic phenotype was mainly driven by features related to regulatory capacity and cofactor metabolism. In particular, the absence of a response regulator and the presence of a lipoate-protein ligase contributed to the prediction of anaerobic potential. Autotrophy was primarily associated with functions linked to hydrogenase activity and maturation. The absence of several hydrogenase-related components consistently reduced the prediction of autotrophy, suggesting that nickel-metallocenter assembly, Hyp maturation machinery, and NiFe/NiFeSe hydrogenases are important indicators of autotrophic potential in this dataset. Predicted cryotolerance was characterized by a combination of functions related to redox and cofactor maintenance, membrane adaptation, compatible-solute synthesis and transport, and genome maintenance. Together, these features point to several complementary strategies that may support survival or activity at low temperatures. Desiccation tolerance was associated with functions involved in oxidative-stress defense, redox buffering, osmoprotection, cell-envelope stabilization, and DNA maintenance. Several of these features indicate that resistance to drying may depend on a broad cellular stress-response network rather than on a single functional system. Halophily was mainly indicated by the absence or reduced contribution of proteins associated with osmoprotection, membrane stability, and redox balance. This pattern suggests that the predictors distinguishing halophilic lifestyles in this dataset may reflect functional differences in how organisms manage hypersaline environments. In contrast, haloresilience was associated with the presence of proteins involved in compatible-solute uptake, ion exchange, sodium-coupled transport, membrane remodeling, redox support, and DNA repair. These features suggest that haloresilient organisms may rely on a broad set of mechanisms that help maintain cellular stability under fluctuating or elevated salt conditions. For psychrophilic lifestyles, the most informative features were largely associated with the absence of proteins involved in proteostasis, membrane fluidity, and translation efficiency. This pattern suggests that reduced representation of these functions was linked to weaker predicted cold-growth performance in the analyzed genomes. Potential indicators of sporulation capability included control proteins, transcriptional regulators, metal-dependent envelope enzymes, signaling molecules, and transport systems. These features are consistent with the complex regulatory and structural changes required for sporulation.

Several proteins appeared among the most predictive features for more than one stress-related trait. For example, COG2130 and COG1292 contributed to both desiccation tolerance and haloresilience. Such proteins may therefore represent broader stress-resistance markers and could be useful candidates for future targeted protein-based screening panels, particularly in the context of polyextremophilic survival strategies.

## Discussion

Icy worlds in the Jovian system (e.g., Europa), ^25^ differ fundamentally from Mars ^1^ as targets for forward contamination: As these environments offer liquid water and relevant key elements (carbon, hydrogen, nitrogen, oxygen, phosphorus, and sulfur) beneath ice shells ^26^, the introduction of a single viable cell or a self-sustaining microbial community into the subsurface could have far-reaching, potentially global consequences ^57^. Hence, these environments could not only permit microbial survival but may also allow proliferation ^58^. At the same time, the decade-long, highly hostile transit environments experienced by microbes on missions such as ESA’s JUICE and NASA’s Europa Clipper space missions impose strong selective filters ^12^. Understanding which microbial contaminants could both survive transit and establish themselves in cryo-subsurface habitats requires functionally oriented risk assessments, not just taxonomic inventories.

Cultivation of candidate contaminants and laboratory phenotyping remain indispensable but are slow and biased against viable but non-culturable (VBNC) organisms ^59^. To complement experimental work, we here developed genome-centric metagenomic approaches coupled with machine learning to predict traits key to multi-extremotolerance (e.g., cryotolerance, desiccation tolerance, halotolerance). This strategy enables trait inference from both 16S rRNA gene amplicon-derived reference genomes and metagenome-assembled genomes (MAGs) ^60^, expanding taxonomic and functional resolution of microbial communities associated with spacecraft assembly cleanrooms ^4^. Two practical applications emerged: (1) mapping 16S rRNA gene amplicon data to relevant reference genomes for cross-referencing with a planetary protection risk catalog, and (2) genome-resolved trait profiling across MAGs to assess community-level risk potential in diverse environments. However, since our research shows that different strains of the same species can exhibit quite distinct traits, our recommendation would be to aim for genome-resolved trait profiling.

Model performance, limitations, and utility depend critically on training data quality and expert curation ^61–66^. Our best-performing classifier (Psychro/Cryotolerance) showed exceptional specificity (1.00) in random-split validation, indicating a conservative tendency to avoid false positives. However, this model was trained and tested on the smallest dataset, limiting confidence in general. By contrast, the desiccation model, while trained on a larger dataset, performed the least well and appears to suffer from heterogeneous or weakly specific training labels. These contrasting outcomes illustrate the trade-off between dataset breadth and trait specificity ^67,68^.

A practical methodological consideration is the balance between prediction confidence and coverage. Applying probability thresholds of ∼0.6–0.7 improved reliability but reduced coverage. For planetary protection, where false negatives carry high consequences, it is appropriate to prioritize conservative thresholds or adopt margin-based approaches that classify uncertain predictions as potential positives. Such conservative strategies reduce the chance of overlooking relevant threats and align with the precautionary principle.

Data curation, expert integration, and the role of the risk catalog expert interpretation are essential to assess model predictions, identify false positives/negatives, and refine training sets. We therefore designed a flexible framework that couples machine learning outputs to a curated planetary protection risk catalog. This catalog links model inferences to expert-validated trait assignments and relevant literature, enabling iterative improvement by incorporating new experimental results and taxonomic updates.

To broaden and strengthen trait classifiers, a systematically structured genome–trait database is a priority. Public, continuously updated datasets would accelerate discovery by enabling: (i) identification of robust genetic markers; (ii) validation and benchmarking of predictive models; and (iii) reproducible, community-wide monitoring. In particular, improving representation of radiation resistance ^17^ and oligotrophy across phylogenetic diversity is urgent because these traits are central to survival under spacelike conditions but currently suffer from sparse, biased datasets. Building balanced collections of resistant and sensitive taxa will require coordinated experimental efforts and time, but it is necessary for robust, generalizable classifiers.

Interpreting prioritized features and biological plausibility, as well as examining which clusters of orthologous groups (COGs) or proteins drive model predictions, yields mechanistic insight and testable hypotheses. Predicting broad phenotypes such as “cold resistance” is inherently more complex than predicting highly conserved traits such as sporulation because multiple evolutionary routes can produce similar phenotypes. Nonetheless, several predicted features in our dataset map to plausible adaptive roles:

Cryotolerance ^21^: Genes involved in NAD biosynthesis (e.g., nicotinate-nucleotide adenylyltransferase) support redox balance at low enzymatic rates; acetyl-CoA hydrolases can buffer acyl-flux and free CoA for membrane lipid remodeling (homeoviscous adaptation) ^69^; ornithine cyclodeaminase generates proline as a cryoprotectant; gyrase-modulating protectors maintain DNA supercoiling homeostasis during temperature shifts ^70^. Conversely, the absence of ribosomal assembly factors (uL29, S19, 23S-binders) or AcpS impairs translation and fatty-acid synthesis required for cold adaptation ^71^.

Desiccation resistance: Catalases, DNA topoisomerases, and DNA glycosylases (MPG family) are critical to manage ROS and repair lesions ^72^ arising during dehydration and rehydration ^15^. BCCT family transporters facilitate import of compatible solutes; D-alanine ligases and teichoic acid transfer strengthen cell envelopes against mechanical stress ^73,74^; cytochrome c and ferredoxin provide electron-dissipation pathways that limit oxidative bursts upon rewetting ^75^.

Halotolerance/Halo-resilience: Transporters and/or synthesis pathways for compatible solutes, antiporters (Na+/H+) that preserve ion homeostasis and pH, ^76,77^ NAD(P)H dehydrogenases supporting motive forces, metal-homeostasis proteins, and photolyases (in high-irradiance saline habitats) collectively mitigate ionic, osmotic, and salt-enhanced oxidative/genotoxic stress ^78,79^.

Sporulation and persistence: Regulatory hubs (e.g., Spo0A pathway) ^80^, metalloproteins for envelope/coating processes, and signaling enzymes (e.g., diguanylate cyclases) appear as key contributors to initiation and coordination of sporulation and long-term survival ^11,81^.

Implications and next steps: The predicted proteins and pathways constitute promising targets for focused experimental screening e.g., multi-primer RT for meta-transcriptomics or targeted qPCR/panel assays, allowing rapid, function-oriented monitoring in cleanrooms and flight hardware. From a planetary protection perspective, integrating machine-predicted trait profiles with curated expert catalogs and targeted experimental validation will provide a more actionable, trait-based risk assessment than taxonomy alone. Finally, while models for cryotolerance, desiccation, and halotolerance show encouraging utility, expanding and balancing training datasets, particularly for radiation resistance and oligotrophy, as well as their experimental validation remains essential. Combining conservative prediction thresholds, margin-based conservative interpretation, and iterative expert curation will help ensure robust, defensible assessments of microbial risk on missions to other habitable worlds in our solar system.

## Conclusions

This study demonstrates the promising application and current limits of using machine learning on genome-centric data to predict phenotypic traits relevant to planetary protection. Models for cryotolerance, haloresilience, and desiccation resistance produced useful, actionable predictions: they can complement taxonomically inferred information, highlight candidate genetic markers, and focus laboratory validation on the organisms and functions most likely to matter for forward contamination. Applying these models to datasets collected around ESA’s JUICE launch site in Kourou shows their practical value for function-based risk assessment and illustrates how computational outputs can be linked to a curated planetary protection risk catalog for expert review.

However, these models are provisional decision-support tools rather than definitive classifiers. Their reliability depends on the specificity and diversity of training data, and some traits, particularly desiccation resistance and radiation resistance, suffer from limited or heterogeneous labels. Consequently, model outputs should be interpreted conservatively and always combined with expert annotation and experimental follow-up. The greatest immediate utility of the models lies in prioritizing targets for validation, thereby allocating laboratory resources to taxa and genes most likely to pose a contamination risk.

In summary, an integrated workflow that connects genome-centric metagenomics, trait-based predictive modeling, and expert-driven experimental validation offers a flexible, probabilistic framework for planetary protection. With continued dataset expansion, methodological refinement, and international collaboration, this approach can strengthen contamination risk assessment and help safeguard extraterrestrial environments during future missions ^32^.

## List of abbreviations (alphabetical order)

AAI: Average Amino-acid Identity
Acc: Accuracy
ANI: Average Nucleotide Identity
COG: Clusters of Orthologous Groups
ddPCR: droplet digital PCR
ECSS: European Cooperation for Space Standardization
ENA: European Nucleotide Archive
ESA: European Space Agency
GTDB: Genome Taxonomy Database
MAG: Metagenome-Assembled Genome
MiDiv: Microbial Diversity project (ESA project name, 2003–2005)
MPG: N-methylpurine DNA glycosylase (DNA repair enzyme family referenced in text)
N50: assembly metric: contig length such that 50% of the assembly is in contigs ≥ this length
NGS: Next-Generation Sequencing
NOB: Nitrite-Oxidizing Bacteria
NTC: No-Template Control
PB/ATP: ATP (adenosine triphosphate): used in viability assays (ATP-based assays)
ProConTra: Probability of Contamination and Transport Mechanisms (project name)
qPCR: quantitative PCR (real-time PCR)
RefSeq: NCBI Reference Sequence database
RNA: Ribonucleic Acid
SHAP: SHapley Additive exPlanations (model-agnostic feature importance method)
SNP: Single-Nucleotide Polymorphism
UV: Ultraviolet (e.g., UV 254 nm sterilization)
VBNC: Viable But Non-Culturable
VBCF: Vienna BioCenter Core Facilities (sequencing facility)

## Glossary of used bioinformatic tools and pipelines

(alphabetical, with short descriptions)

BBMap: High-performance read mapper and suite for read processing and filtering.
Bowtie2: Fast aligner for mapping sequencing reads to reference sequences (short reads).
CheckM2: Genome quality assessment tool estimating completeness and contamination of genome bins.
DASTool: Bin refinement tool that integrates multiple binning results to improve MAG quality.
DIAMOND: Fast protein aligner for sensitive sequence similarity searches against large protein databases.
DRAM: Distilled and Refined Annotation of Metabolism. Functional annotation pipeline focused on metabolic reconstruction and genome annotation.
dRep: Dereplication tool to cluster and select representative genomes from genome sets.
eggNOG mapper: Functional annotation tool mapping genes to orthology groups and functional categories (eggNOG).
GRiD: Growth Rate InDex; infers replication activity from coverage patterns (peak-to-trough ratios).
GTDBtk: GTDB toolkit for taxonomic classification using the GTDB database.
inStrain: Strain-level genomic comparison and microdiversity profiling from mapped metagenomic reads.
Kraken2: Taxonomic classifier that assigns reads to taxa using exact k-mer matches (fast profiling).
Linclust: Ultrafast clustering of protein sequences (used to cluster redundant genes).
metaSPAdes: Metagenome assembler optimized for complex metagenomic datasets.
MetaBAT2: Automated binning tool that groups contigs into genome bins using coverage and composition.
Minimap2: Fast sequence aligner suitable for mapping long and short reads to references.
MMseqs2: Sensitive and fast sequence search and clustering toolkit for large sequence sets.
Phenotrex: phenotype prediction package; used to run phenotype models.
Prodigal: Gene prediction program for prokaryotic genomes (calling coding sequences).
Snakemake: Workflow management system used to organize and reproduce bioinformatic pipelines.

## Declarations

### Ethics approval and consent to participate

NA

### Consent for publication

All authors approved the manuscript and give their consent for publication.

### Availability of data and material

All raw sequencing data were submitted to ENA (https://www.ebi.ac.uk/ena/browser/home) and are accessible under the following project: PRJEB104687 (ERP185943, Jupiter Icy Moons Explorer).

Data for our meta-analysis were retrieved from the following repositories: Representative MAGs from the International Space Station: NASA’s Open Science Data Repository (OSDR), OSD-730, OSD-731, OSD-732, OSD-733; and NCBI SRA database (Project: PRJNA1328788).

Representative MAGs from cleanroom meta-study: The Open University. Dataset. (doi:10.21954/ou.rd.29383073)

Genome bins from deeply sequenced healthy skin microbiome: NCBI SRA database (Project: PRJNA1108908).

Selected genomes of ESA’s strain collection at DSMZ https://www.dsmz.de/ Reference collection of genomes and isolates of the skin microbiome: https://ftp.ebi.ac.uk/pub/databases/metagenomics/genome_sets/skin_microbiome/

### Code availability

The present study did not generate new code, and the mentioned tools used for the data analysis were applied with default parameters unless specified otherwise in the Methods and our GitHub repo: https://github.com/Mechah/ProConTra

### Competing interests

The authors declare that they have no competing interests.

### Funding

This study was funded by ESA Contract No. 4000141696/23/NL/AR/ahh coordinated by PR, ESA Cooperative Agreement No. 4000148372/25/NL/GLC given to AM, and ESA Contract No. 4000136034/21/NL/AR/zk coordinated by PR.

### Authors’ contributions

Study concept: PR, SS, CM-E, AM, JC, MAS; Sampling: CK, AM; Machine learning: TM; Data analysis: TM, AM; Data interpretation: TM, AM, PR; Wrote the manuscript: AM; All authors approved the manuscript.

## Acknowledgements

We greatly value the computational resources of the MedBioNode provided by Medical University of Graz ZMF Galaxy Team: Core Facility Computational Bioanalytics, Medical University of Graz, funded by the Austrian Federal Ministry of Education, Science and Research, Hochschulraum-Strukturmittel 2016 grant as part of BioTechMed Graz. In addition, we would like to thank for additional computing capacity at the Life Science Compute Cluster (LiSC) and support by Thomas Rattei and the Computational Systems Biology group at the University of Vienna.

We would like to thank Jack A. Gilbert and Megan Hill (UCSD, USA), Karen Olsson-Francis and Michael Macey (OU, UK), and Ruediger Pukall (DSMZ, GER) for providing relevant data for our meta-analysis of phenotypic traits.

**Large Language Models (LLMs)** were used exclusively for linguistic revisions and text summarization, but not for content generation.

## Notes

https://github.com/Mechah/ProConTra

### Competing Interest Statement

The authors have declared no competing interest.

