## Supplementary Information for "Functional profiling of spacecraft cleanroom microbiomes through genome-wide phenotype predictions"

#### Affiliations

### Supplementary Methods and Results

#### 1. Data acquisition and processing

1.1 Positive (trait-positive) dataset

Species and strain selection primarily used the Omnicrobe 1.2 database [1] to identify candidate organisms associated with target biotopes/phenotypes (e.g., halotolerance, halophily, hypersaline lake environments). For each selected species we retrieved RefSeq assemblies (protein .faa files) using the RefSeq bacteria assembly summary (https://ftp.ncbi.nlm.nih.gov/genomes/refseq/bacteria/assembly_summary.txt). These assemblies and their metadata (assembly level, taxIDs, taxonomy, source) were used to generate the final positive sets (https://github.com/Mechah/ProConTra/tree/main/Model_datasets).

1.2 Negative (closely related control) dataset

Negative candidates were initially identified by extracting 16S rRNA gene sequences for the positive species via NCBI Entrez (Biopython v1.75) and performing BLASTn (BLAST 2.12.0+) against a 16S rRNA gene reference database (BioProject PRJNA33175) to obtain closely related taxa. Candidate negatives were then filtered to remove species associated with unwanted habitat/biotope keywords using Omnicrobe 1.2 OntoBiotope IDs (e.g., salt, halotolerance, saline water) and further refined using BacDive API (v0.2) metadata (growth at NaCl concentrations, halophily, temperature range). Any culture reporting growth above 3% NaCl was excluded from haloresilient negative sets. Potential negatives were ranked by BLAST bitscore; selected and excluded candidates are annotated in the Negative_set parameters (Supplementary Table S1). The final paired positive/negative lists are provided in Model_Datasets (https://github.com/Mechah/ProConTra/tree/main/Model_datasets).

**Supplementary Table 1**: Model details including positive and negative cut-offs and OntoBiotope IDs used for biotope/habitat definitions and negative set filtering. Species found to be associated with a certain habitat or more specifically ID were eliminated as a potential negative candidate. The selection parameters for each model were carefully chosen, particularly for use during the refinement stage. The parameters were designed to create a distinct separation between the positive and negative sets, based on specific environmental conditions and growth capabilities. This was done to ensure that the organism had distinct traits that the algorithm could effectively differentiate.

| Model | Haloresilient2 | Cryotolerance5 | Desiccation2 |
|---|---|---|---|
| Genomes | 560 | 184 | 638 |
| Positive_set parameters | Hypersaline Habitat<br>Halotolerant/phile<br>growth above 10% NaCl | Glacier Habitat<br>Psychrophile<br>Growth below 4°C | Desert Habitat |
| Negative_set parameters | Growth only below 3% NaCl | Growth only above 10°C | Not Desert Habitat |
| OntoBiotope_IDs | OBT:000338<br>OBT:003226<br>OBT:002940<br>OBT:003261<br>OBT:001290<br>OBT:002720<br>OBT:003260<br>OBT:003333 | OBT:000681<br>OBT:002709<br>OBT:002551 | OBT:000592 |
| Definition | salt<br>saline lake<br>soda lake<br>hypersaline lake<br>saline wetland<br>hypersaline water<br>highly alkaline saline soda lake<br>alkaline lake | ice<br>glacier<br>permafrost | desert |

### 2. Feature generation and genotype/phenotype files

Protein .faa files from RefSeq assemblies were annotated to COGs (eggNOG v5) using PyHMMER [2]. Phenotrex v0.6.0 [3] was used to generate genotype files (genome_assembly_id mapped to orthologous groups with alignment-score > 0.5) and phenotype files (assembly IDs with binary trait labels) following the Phenotrex usage protocol (https://phenotrex.readthedocs.io/en/latest/usage.html). COG presence/absence (and counts where applicable) constituted the feature representation for model training.

### 3. Model development and refinement

#### 3.1 Model choice and training pipeline

We trained Extreme Gradient Boosting (XGBoost) [4] classifiers through Phenotrex to predict microbial traits relevant to planetary protection (PP). Hyperparameter tuning used Phenotrex's optimization routine (–optimize) with 100 optimization iterations (–optimize_n_iter) and outputs saved (–optimize_out). Optimized parameters (n_estimators, max_depth, gamma, min_child_weight, colsample_bytree, subsample) were then applied during final training. Individual model hyperparameters are listed in Table 1.

3.2 Cross-validation and iterative data curation
To identify and remove mislabeled or ambiguous assemblies (false positives from text-mined sources), we applied repeated cross-validation (5 folds × 10 replicates). Genomes with consistently poor cross-validation performance (high misclassification probability) were manually reviewed against literature; assemblies confirmed as mislabeled were removed (documented in the repository [https://github.com/Mechah/ProConTra/tree/main/Refinements] with citations). This iterative refinement was applied during dataset fusions (e.g., progressive fusions for Haloresilience2) and is summarized in the Refinement section (Supplementary Results).

4. Model evaluation

4.1 Evaluation strategies
We evaluated models using two complementary splits:

Random split: standard 80:20 train:test random partition (sklearn.model_selection.train_test_split).

Genomic-distance split (AAI-based): to test generalization to taxonomically distant genomes, we computed pairwise Average Amino Acid Identity (AAI) between .faa files using CompareM (v0.1.2). Test sets were formed by selecting genomes with the lowest summed AAI distances (top 20% lowest-sum genomes) to ensure the test set comprises the most distant genomes relative to the training set.

4.2 Performance metrics and thresholds
We report accuracy, sensitivity (true positive rate), specificity (true negative rate), and balanced accuracy (average of sensitivity and specificity). Models provide a probabilistic confidence per prediction; reported results use multiple confidence thresholds (e.g., 0.5, 0.6, 0.7) to characterize trade-offs between sensitivity and specificity. Histogram and boxplot visualizations of confidence distributions and accuracy at thresholds are provided in Supplementary Figures 1 and 2.

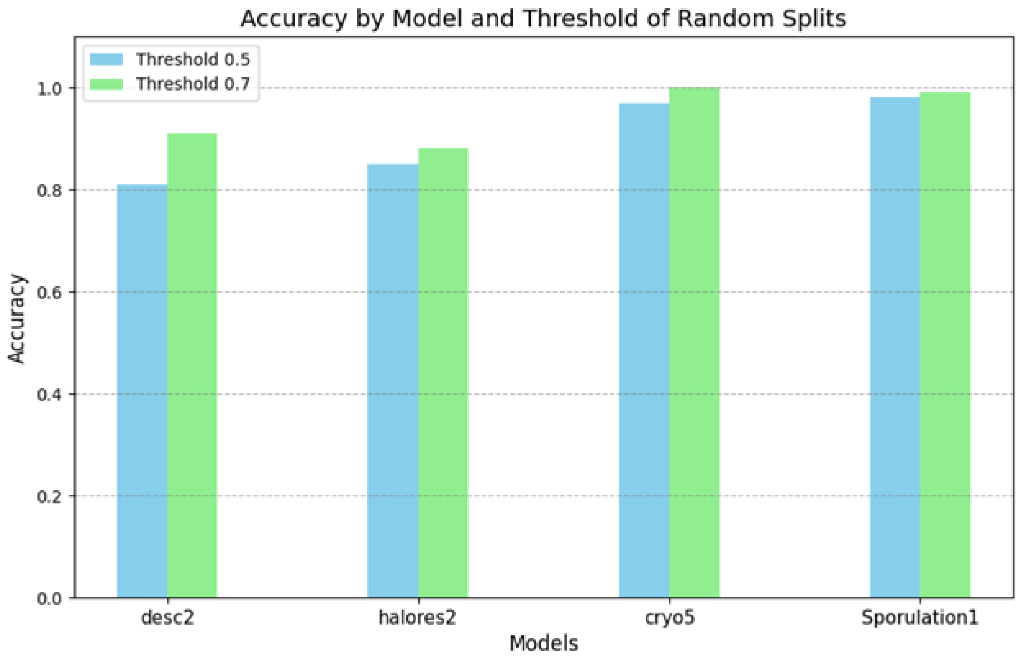


**Supplementary Figure 1**: Random Split Histogram: Histogram showing model accuracies at confidence thresholds of 0.5 and 0.6 for the random split datasets

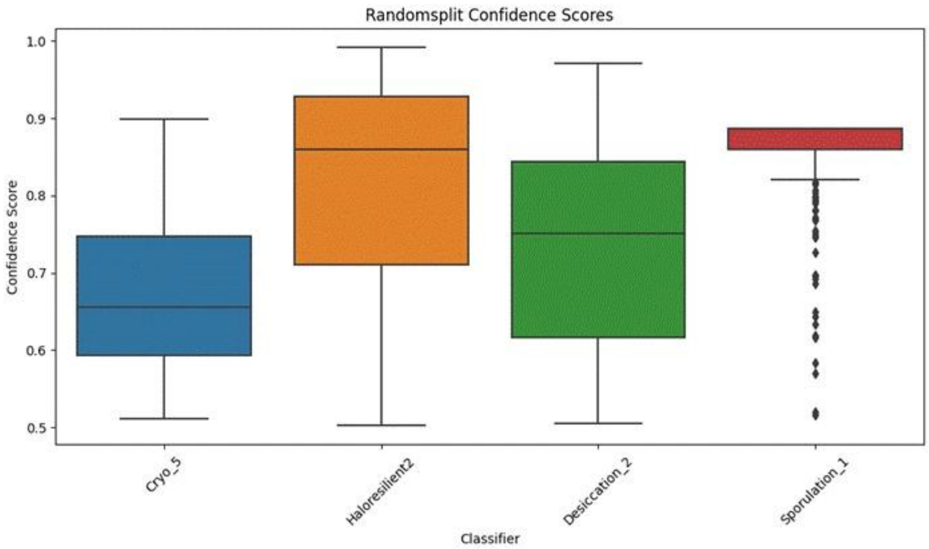


**Supplementary Figure 2**: Box plot depicting the distribution of confidence scores for each classifier. The spread and median values indicate the model's certainty in its predictive capabilities.

4.3 Observations from AAI vs random splits

As expected, AAI-based splits produced lower accuracies than random splits due to increased taxonomic distance; model ranking remained consistent. The AAI split caused greater loss of high-confidence predictions at stricter thresholds (e.g., ~64%

lost predictions at 0.7 threshold on average vs ~41.6% for random split). Detailed per-model performance and threshold effects are in Table 2 and Supplementary Figure 3.

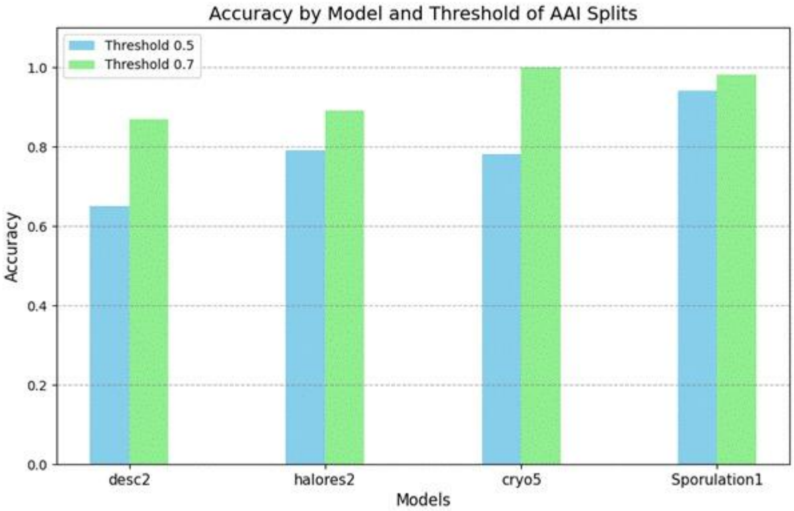


**Supplementary Figure 3**: Histogram showing model accuracies at confidence thresholds of 0.5 and 0.6 for the AAI split datasets.

5. Completeness and contamination simulations

Using Phenotrex's cccv resampling, we simulated incomplete and contaminated genomes by randomly removing COGs (to mimic incompleteness) and injecting COGs from genomes of the opposite class (to mimic contamination). Cross-validation under these perturbations shows that models remain stable down to ~80% completeness and up to ~20% contamination for many classifiers. However, new trait models are more sensitive to contamination than established models (e.g., sporulation classifier), with accuracy declining more rapidly as contamination increases. Results are plotted in Supplementary Figures 4 and 5 (Completeness vs Mean Balanced Accuracy; Contamination vs Mean Balanced Accuracy).

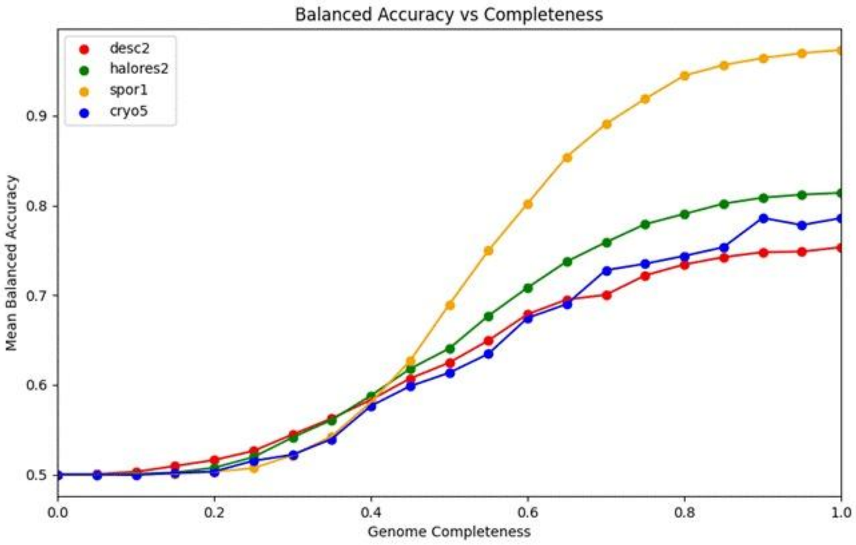


**Supplementary Figure 4**: Completeness vs Mean Balanced Accuracy: Analysis of genome completeness based on cross-validation simulations.

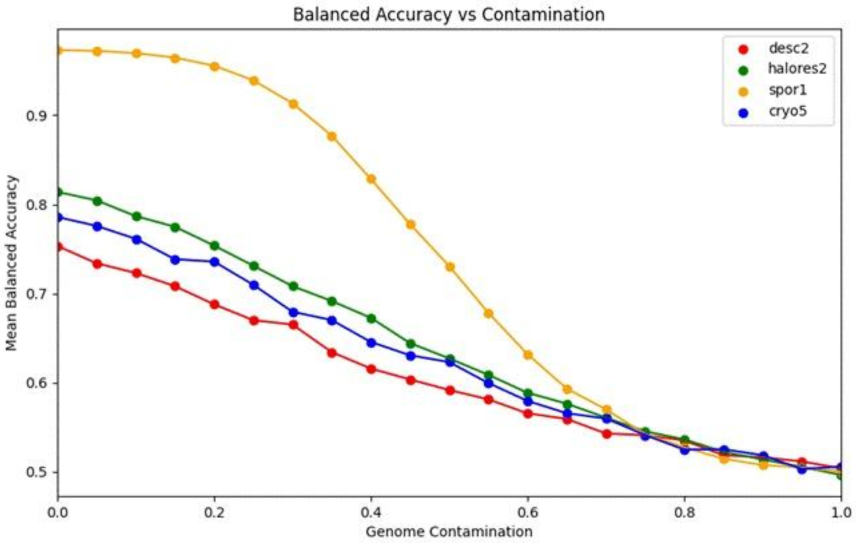


**Supplementary Figure 5**: Contamination vs Mean Balanced Accuracy: Analysis of genome contamination based on cross-validation simulations.

### 6. Cleanroom isolate analysis (PP-VERI dataset)

#### 6.1 Data and limitations

A datasheet from the PP-VERI project (Medical University of Graz) listing cultivable isolates from planetary protection campaigns was expanded with additional cleanroom isolates sourced via BacDive (https://bacdive.dsmz.de). This dataset is biased toward culturable, assay-selected organisms (not a comprehensive representation of cleanroom bioburden) and should be interpreted accordingly.

#### 6.2 RefSeq genome selection and trait predictions

RefSeq genomes matching the curated isolate list (n = 126) were analyzed with the new trait models plus existing phenotypes (sporulation, anaerobe) available via PhenDB. Phenotrex outputs a probability/confidence in (0.5, 1) for positive predictions and (0, 0.5) for negatives; for conservative screening we explored lowering the YES/NO boundary to 0.4 to capture borderline positives. Trait abundances and confidence distributions are summarized (Supplementary Figures 6 and 7). Bacillales comprised ~40% of the RefSeq genomes—likely due to the ECSS assay selection for heat-resistant spore formers.

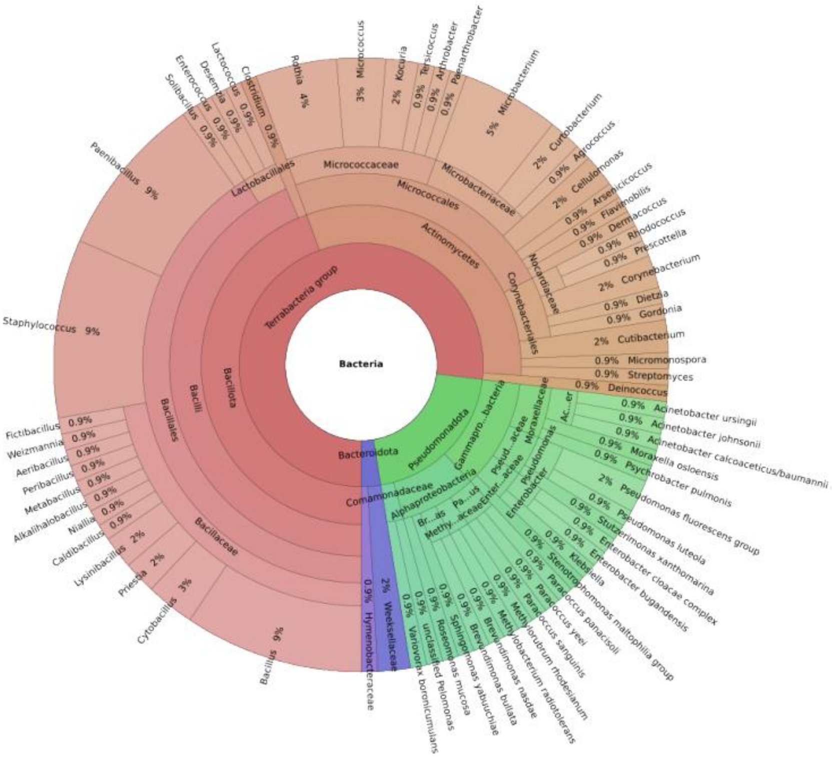


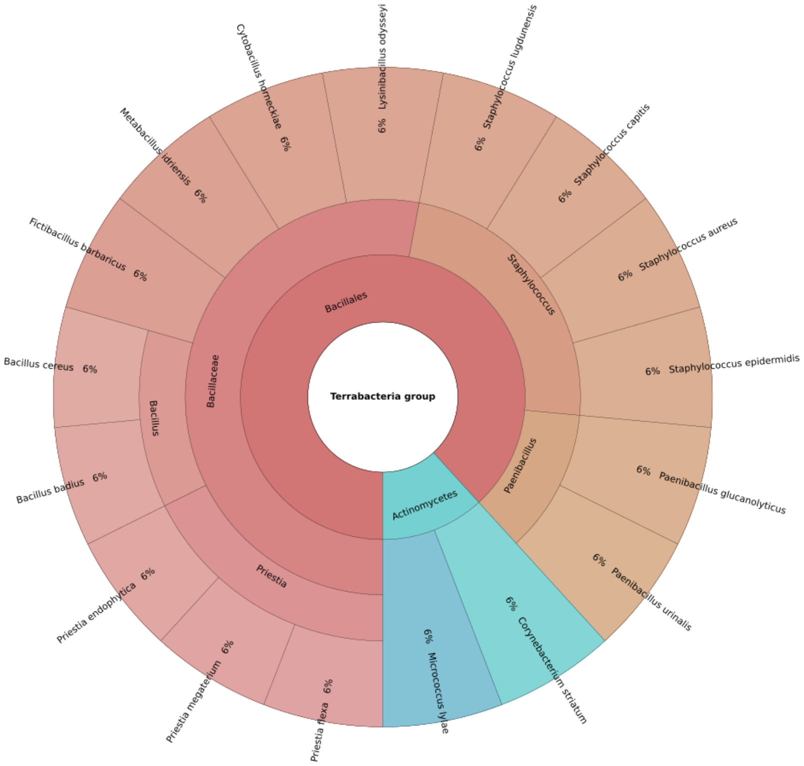


**Supplementary Figure 6**: Krona chart of bacterial cleanroom isolates: The plot provides an overview of bacteria which were isolated from cleanrooms (top: all taxa, bottom: selected Bacteria with planetary protection relevance according to predicted traits).

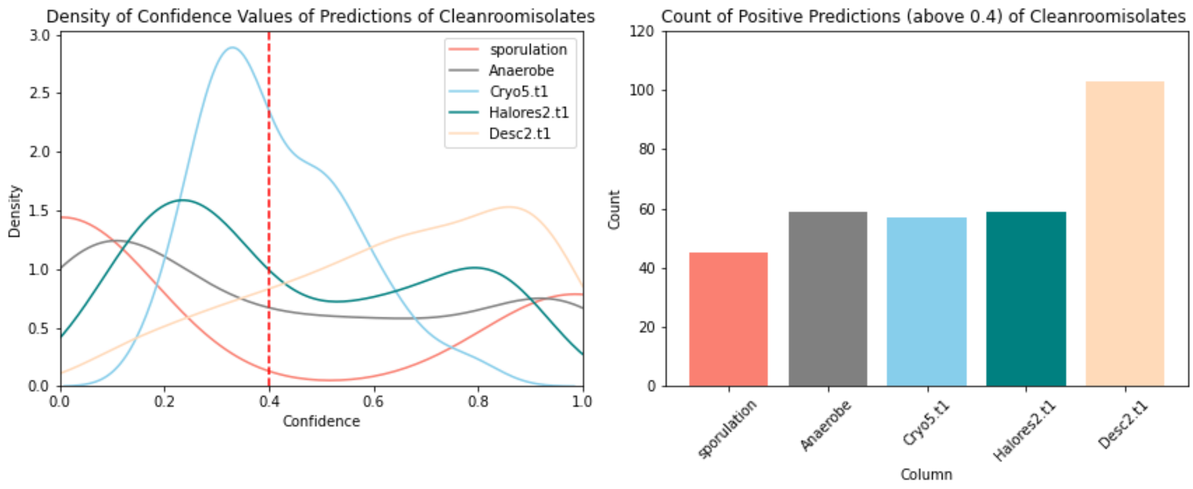


**Supplementary Figure 7**: Confidence and trait abundance in cleanroom set: The density of confidence values are illustrated where 0 to 0.4 are now considered as 'NO' and above 0.4 considered 'YES'. The histogram shows the sum count of predicted microbial traits (above 0.4) relevant to Planetary Protection within the cleanroom dataset.

6.3 Candidate-risk organisms

Applying selection criteria (organisms predicted YES for sporulation or anaerobic capability, and additional survival-related traits), we identified 17 species of interest for PP (e.g., *Paenibacillus glucanolyticus, Bacillus badius, Paenibacillus urinalis, Fictibacillus barbaricus, Lysinibacillus odysseyi*). *Paenibacillus glucanolyticus* exhibited strong, high-confidence YES predictions across all models. The candidate list (with per-species prediction confidences) is provided in Supplementary Figure 8; these candidates require validation with genome data from the isolates and laboratory assays before operational use in risk catalogues.

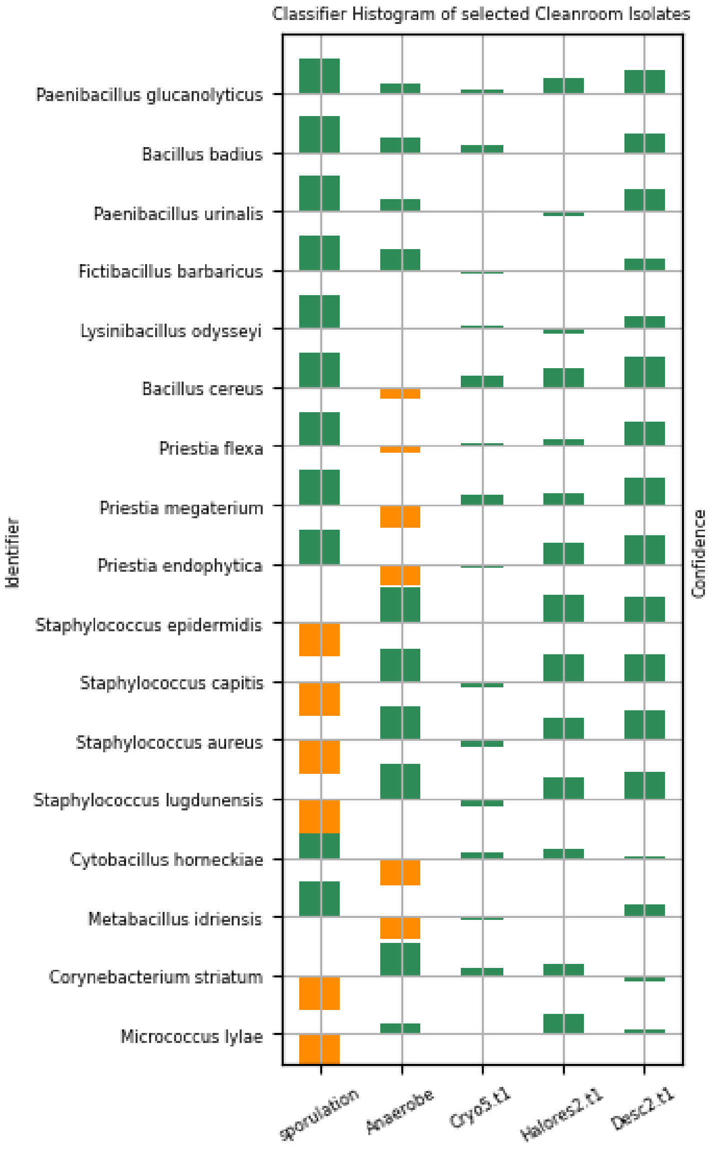


**Supplementary Figure 8**: Selected microbial isolates ranked by positive hit count: This figure illustrates the predictions of microbial isolates with 4 or 5 positive predictions (YES). If the prediction is below 0.5 it is shown as below the section. Green indicates it is within the 0.4 margin (YES), while orange indicates it is below the margin.

### 6.4 Taxonomic profile extension

We also used 16S rRNA gene amplicon taxonomic profiles from ESA PP-VERI campaigns 17–33 to select representative reference genomes for phenotype prediction; 17 of 127 species showed >=3/6 positive traits relevant to icy-moon survivability. *Bacillus cereus* was notable for 4 positive traits (desiccation resistance, sporulation, haloresilience, cryotolerance).

7. Scripts, reproducibility and resources

All scripts and pipeline configurations used for data acquisition, filtering, feature generation, model training, cross-validation, AAI computations, and simulations are provided at https://github.com/Mechah/ProConTra/tree/main/Scripts. Refinement decisions and literature citations for removed assemblies are documented in the repository's refinement logs.

8. Summary of key formulas and metrics

- TP, TN, FP, FN definitions as used in metric calculations.

- Accuracy = (TP + TN) / (TP + TN + FP + FN)

- Sensitivity (TPR) = TP / (TP + FN)

- Specificity (TNR) = TN / (TN + FP)

- Balanced accuracy = (Sensitivity + Specificity) / 2

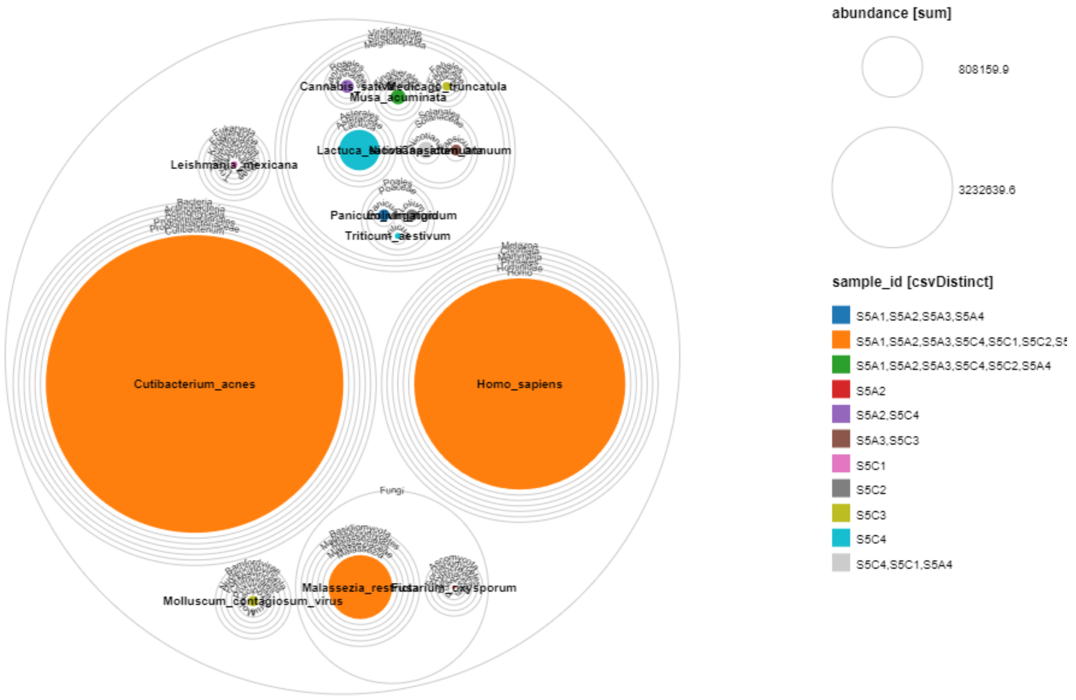


**Supplementary Figure 9**: Read based profiling of all domains of life in Kourou during the JUICE mission. Color code refers to the prevalence in specific samples.

**Supplementary Table 2**: Overall comparative statistics between traits and studies.

| trait | prediction | statistic | df | p | n_groups | n_total | p_fmt | signif |
|---|---|---|---|---|---|---|---|---|
| Anaerobe | NO | 68.60979 | 6 | 7.88E-13 | 7 | 554 | 7.88e-13 | *** |
| Anaerobe | YES | 114.1499 | 6 | 2.75E-22 | 7 | 1314 | <2e-16 | *** |
| AUTO | NO | 124.2314 | 6 | 2.1E-24 | 7 | 1591 | <2e-16 | *** |
| AUTO | YES | 22.36781 | 6 | 0.00104 | 7 | 277 | 0.00104 | ** |
| Cryotolerant.t1 | NO | 76.59372 | 6 | 1.8E-14 | 7 | 1658 | 1.8e-14 | *** |
| Cryotolerant.t1 | YES | 7.443919 | 6 | 0.282 | 7 | 210 | 0.282 | |
| Desiccationtolerance.t1 | NO | 11.59115 | 6 | 0.0717 | 7 | 1209 | 0.0717 | . |
| Desiccationtolerance.t1 | YES | 80.48727 | 6 | 2.83E-15 | 7 | 659 | 2.83e-15 | *** |
| HALO | NO | 104.8434 | 6 | 2.44E-20 | 7 | 1816 | <2e-16 | *** |
| HALO | YES | 22.6789 | 4 | 0.000147 | 5 | 52 | 0.000147 | *** |
| Haloresilience.t1 | NO | 75.75543 | 6 | 2.68E-14 | 7 | 1368 | 2.68e-14 | *** |
| Haloresilience.t1 | YES | 36.79461 | 6 | 1.93E-06 | 7 | 500 | 1.93e-06 | *** |
| PSYCHRO | NO | 143.9295 | 6 | 1.48E-28 | 7 | 1856 | <2e-16 | *** |
| PSYCHRO | YES | 3.703297 | 5 | 0.593 | 6 | 12 | 0.593 | |
| sporulation | NO | 233.4442 | 6 | 1.41E-47 | 7 | 1794 | <2e-16 | *** |
| sporulation | YES | 12.06355 | 4 | 0.0169 | 5 | 74 | 0.0169 | * |

**Supplementary Table 3**: Pairwise comparative statistics between traits and selected studies. Please check GitHub repository due to size limitations: https://github.com/Mechah/ProConTra

**Supplementary Table 4**: Pairwise comparative statistics between traits predicted for JUICE Kourou and other studies. Please check GitHub repository due to size limitations: https://github.com/Mechah/ProConTra

**Supplementary Table 5**: Pairwise comparative statistics between traits of *Staphylococcus* predicted for JUICE Kourou and other studies.

| trait | study_group | total | yes | percent_yes | overall_test | overall_pvalue |
|---|---|---|---|---|---|---|
| Anaerobe | JUICE_Kourou | 25 | 25 | 100 | chisq.test (low expected) | NA |
| Anaerobe | DSMZ_ESA_strains | 34 | 34 | 100 | chisq.test (low expected) | NA |
| Anaerobe | reference datasets | 183 | 183 | 100 | chisq.test (low expected) | NA |
| AUTO | JUICE_Kourou | 25 | 2 | 8 | chisq.test (low expected) | 0.18719227511871023 |
| AUTO | DSMZ_ESA_strains | 34 | 0 | 0 | chisq.test (low expected) | 0.18719227511871023 |
| AUTO | reference datasets | 183 | 5 | 2.73224043715847 | chisq.test (low expected) | 0.18719227511871023 |
| Cryotolerant.t1 | JUICE_Kourou | 25 | 2 | 8 | chisq.test (low expected) | 0.5742477726951674 |
| Cryotolerant.t1 | DSMZ_ESA_strains | 34 | 1 | 2.9411764705882355 | chisq.test (low expected) | 0.5742477726951674 |
| Cryotolerant.t1 | reference datasets | 183 | 7 | 3.8251366120218577 | chisq.test (low expected) | 0.5742477726951674 |
| Desiccationtolerant.t1 | JUICE_Kourou | 0 | 0 | NA | NA | NA |
| Desiccationtolerant.t1 | DSMZ_ESA_strains | 0 | 0 | NA | NA | NA |
| Desiccationtolerant.t1 | reference datasets | 0 | 0 | NA | NA | NA |
| HALO | JUICE_Kourou | 25 | 1 | 4 | chisq.test (low expected) | 0.01280386374208497 |
| HALO | DSMZ_ESA_strains | 34 | 0 | 0 | chisq.test (low expected) | 0.01280386374208497 |
| HALO | reference datasets | 183 | 0 | 0 | chisq.test (low expected) | 0.01280386374208497 |
| Haloresilience.t1 | JUICE_Kourou | 25 | 19 | 76 | chisq.test (low expected) | 0.01765387985876927 |
| Haloresilience.t1 | DSMZ_ESA_strains | 34 | 34 | 100 | chisq.test (low expected) | 0.01765387985876927 |
| Haloresilience.t1 | reference datasets | 183 | 151 | 82.51366120218579 | chisq.test (low expected) | 0.01765387985876927 |
| PSYCHRO | JUICE_Kourou | 25 | 0 | 0 | chisq.test (low expected) | NA |
| PSYCHRO | DSMZ_ESA_strains | 34 | 0 | 0 | chisq.test (low expected) | NA |
| PSYCHRO | reference datasets | 183 | 0 | 0 | chisq.test (low expected) | NA |
| sporulation | JUICE_Kourou | 25 | 0 | 0 | chisq.test (low expected) | NA |
| sporulation | DSMZ_ESA_strains | 34 | 0 | 0 | chisq.test (low expected) | NA |
| sporulation | reference datasets | 183 | 0 | 0 | chisq.test (low expected) | NA |

**Supplementary Table 6**: Overall summary of comparative statistics between traits of *Staphylococcus* predicted for JUICE Kourou and other studies.

| trait | total_across_groups | yes_across_groups | percent_yes_overall | overall_test | overall_pvalue |
|---|---|---|---|---|---|
| AUTO | 242 | 7 | 2.8925619834710745 | chisq.test (low expectec | 0.18719227511871023 |
| Anaerobe | 242 | 242 | 100 | chisq.test (low expectec | NA |
| Cryotolerant.t1 | 242 | 10 | 4.132231404958677 | chisq.test (low expectec | 0.5742477726951674 |
| Desiccationtolerant.t: | 0 | 0 | NA | NA | NA |
| HALO | 242 | 1 | 0.4132231404958678 | chisq.test (low expectec | 0.01280386374208497 |
| Haloresilience.t1 | 242 | 204 | 84.29752066115702 | chisq.test (low expectec | 0.01765387985876927 |
| PSYCHRO | 242 | 0 | 0 | chisq.test (low expectec | NA |
| sporulation | 242 | 0 | 0 | chisq.test (low expectec | NA |

**Supplementary Table 7**: Exploratory statistics of the conducted meta-analysis.

Confidence by study summary

| study | variable | N_nonmiss | median | p25 | p75 |
|---|---|---|---|---|---|
| ISS_MT123 | confidence | 32592 | 89715 | 7259 | 9706 |
| SMGC | confidence | 26124 | 9368 | 825375 | 9.7813E+15 |
| JUICE_Kourou | confidence | 8904 | 85145 | 6904 | 948325 |
| DSMZ_ESA_strains | confidence | 5460 | 9577 | 867375 | 985525 |
| AoHS | confidence | 2016 | 9428 | 843075 | 9812 |
| ensemble_binning | confidence | 1848 | 91055 | 7.6798E+15 | 9699 |
| ISS_MT3 | confidence | 1512 | 95145 | 8.599E+15 | 9.8323E+15 |

Overall confidence summary

| variable | N_nonmiss | median | p25 | p75 |
|---|---|---|---|---|
| confidence | 78456 | 9157 | 7681 | 9743 |

Identifiers by study and prediction

| study | prediction | n_rows | n_unique_identifiers |
|---|---|---|---|
| AoHS | NO | 1641 | 48 |
| AoHS | YES | 375 | 48 |
| DSMZ_ESA_strains | NO | 4399 | 130 |
| DSMZ_ESA_strains | YES | 1061 | 130 |
| ensemble_binning | NO | 1519 | 44 |
| ensemble_binning | YES | 329 | 44 |
| ISS_MT123 | NO | 25836 | 776 |
| ISS_MT123 | YES | 6756 | 776 |
| ISS_MT3 | NO | 1220 | 36 |
| ISS_MT3 | YES | 292 | 36 |
| JUICE_Kourou | NO | 7145 | 212 |
| JUICE_Kourou | YES | 1759 | 212 |
| SMGC | NO | 21254 | 622 |
| SMGC | YES | 4870 | 622 |

Identifiers counts by study

| study | n_rows | n_unique_id | n_missing_identifier |
|---|---|---|---|
| ISS_MT123 | 32592 | 776 | 0 |
| SMGC | 26124 | 622 | 0 |
| JUICE_Kourou | 8904 | 212 | 0 |
| DSMZ_ESA_strains | 5460 | 130 | 0 |
| AoHS | 2016 | 48 | 0 |
| ensemble_binning | 1848 | 44 | 0 |
| ISS_MT3 | 1512 | 36 | 0 |

Predictions by study summary

| study | prediction | n | pct |
|---|---|---|---|
| AoHS | NO | 1641 | 8.13988E+15 |
| AoHS | YES | 375 | 1.86012E+16 |
| DSMZ_ESA_strains | NO | 4399 | 8.05678E+15 |
| DSMZ_ESA_strains | YES | 1061 | 1.94322E+15 |
| ensemble_binning | NO | 1519 | 8.2197E+14 |
| ensemble_binning | YES | 329 | 1.7803E+16 |
| ISS_MT123 | NO | 25836 | 7.9271E+15 |
| ISS_MT123 | YES | 6756 | 2.0729E+15 |
| ISS_MT3 | NO | 1220 | 8.06878E+15 |
| ISS_MT3 | YES | 292 | 1.93122E+16 |
| JUICE_Kourou | NO | 7145 | 8.02448E+15 |
| JUICE_Kourou | YES | 1759 | 1.97552E+16 |
| SMGC | NO | 21254 | 8.13581E+15 |
| SMGC | YES | 4870 | 1.86419E+16 |

Overall prediction summary

| prediction | n | pct |
|---|---|---|
| NO | 63014 | 8.0318E+15 |
| YES | 15442 | 1.9682E+16 |

P-values per tested variable

| variable | test | p.value |
|---|---|---|
| confidence | kruskal.test | 0 |
| prediction | chisq.test | 238335777 |
| trait | chisq.test | 1 |
| type | chisq.test | 0 |
| domain | chisq.test | 0 |
| phylum | chisq.test | 0 |

### **Supplementary Table 8**: Feature importance (top 10) for JUICE Korou trait: anaerobe

| | | Anaerobe | | | |
|---|---|---|---|---|---|
| **Feature** | **Mean SHAP If Present** | **Mean SHAP If Absent** | **N(present)** | **N(absent)** | **Feature Annotation** |
| COG3011 | -0.02887 | 0.16753 | 8 | 17 | Protein conserved in bacteria |
| COG3707 | 0 | 0.21747 | 11 | 14 | response regulator |
| COG2095 | 0 | 0.10226 | 6 | 19 | MarC family integral membrane protein |
| COG1004 | 0 | 0.15848 | 14 | 11 | Belongs to the UDP-glucose GDP-mannose dehydrogenase family |
| COG2096 | 0 | 0.07354 | 3 | 22 | Adenosyltransferase |
| COG3655 | 0 | 0.09977 | 10 | 15 | Transcriptional regulator |
| COG0010 | 0 | 0.10358 | 11 | 14 | Belongs to the arginase family |
| COG2355 | 0 | 0.05501 | 2 | 23 | Zn-dependent dipeptidase, microsomal dipeptidase |
| COG0095 | 0.0854 | 0 | 14 | 11 | Lipoate-protein ligase |
| COG5485 | 0 | 0.05497 | 4 | 21 | Ester cyclase |

### **Supplementary Table 9**: Feature importance (top 10) for JUICE Korou trait: autotroph

| | | | | | AUTO |
|---|---|---|---|---|---|
| **Feature** | **Mean SHAP If Present** | **Mean SHAP If Absent** | **N(present)** | **N(absent)** | **Feature Annotation** |
| COG0375 | 0 | -0.15754 | 1 | 24 | protein maturation |
| COG0378 | 0 | -0.20163 | 7 | 18 | Facilitates the functional incorporation of the urease nickel metallocenter. This process requires GTP hydrolysis, probably effectuated by UreG |
| COG2862 | 0 | -0.14628 | 5 | 20 | Uncharacterized protein family, UPF0114 |
| COG0409 | 0 | -0.11151 | 1 | 24 | Hydrogenase expression formation protein |
| COG0068 | 0 | -0.1022 | 1 | 24 | Along with HypE, it catalyzes the synthesis of the CN ligands of the active site iron of NiFe -hydrogenases using carbamoylphosphate as a substrate. |
| COG0374 | 0 | -0.09792 | 1 | 24 | Belongs to the NiFe NiFeSe hydrogenase large subunit family |
| COG0298 | 0 | -0.08914 | 1 | 24 | carbon dioxide binding |
| COG1740 | 0 | -0.08159 | 1 | 24 | oxidoreductase activity, acting on hydrogen as donor, iron-sulfur protein as acceptor |
| COG0247 | 0 | -0.15908 | 14 | 11 | lactate metabolic process |
| COG1566 | 0 | -0.08302 | 9 | 16 | PFAM secretion protein HlyD family protein |

### **Supplementary Table 10**: Feature importance (top 10) for JUICE Korou trait: cryotolerance

| | | | Cryotolerance | | |
|---|---|---|---|---|---|
| **Feature** | **Mean SHAP If Present** | **Mean SHAP If Absent** | **N(present)** | **N(absent)** | **Feature Annotation** |
| COG0071 | -0.11156 | 0.26611 | 10 | 15 | Belongs to the small heat shock protein (HSP20) family |
| COG2509 | 0.26058 | -0.18577 | 1 | 24 | 5-formyltetrahydrofolate cyclo-ligase activity |
| COG0427 | 0.14943 | -0.1435 | 9 | 16 | acetyl-CoA hydrolase |
| COG3177 | 0.15192 | -0.12428 | 10 | 15 | Filamentation induced by cAMP protein fic |
| COG1358 | 0.22014 | -0.104 | 3 | 22 | cellular component organization or biogenesis |
| COG3554 | 0.16976 | -0.0811 | 4 | 21 | Major facilitator Superfamily |
| COG2423 | 0.08568 | -0.08477 | 10 | 15 | ornithine cyclodeaminase activity |
| COG4135 | | -0.07138 | 0 | 25 | transport system, permease component |
| COG4111 | 0.06946 | -0.0706 | 9 | 16 | nicotinate-nucleotide adenylyltransferase activity |
| COG3449 | 0.07942 | -0.06232 | 2 | 23 | Inhibits the supercoiling activity of DNA gyrase. Acts by inhibiting DNA gyrase |

### **Supplementary Table 11**: Feature importance (top 10) for JUICE Korou trait: desiccationtolerance

**Desiccationtolerance**

| Feature | Mean SHAP If Present | Mean SHAP If Absent | N(present) | N(absent) | Feature Annotation |
|---|---|---|---|---|---|
| COG2010 | -0.04091 | 0.39987 | 12 | 13 | Cytochrome c |
| COG3546 | 0.18078 | -0.2319 | 5 | 20 | catalase activity |
| COG2130 | 0.11797 | -0.12823 | 9 | 16 | NADP-dependent |
| COG1054 | 0.09979 | -0.14938 | 13 | 12 | Belongs to the UPF0176 family |
| COG1292 | 0.06645 | -0.13912 | 13 | 12 | Belongs to the BCCT transporter (TC 2.A.15) family |
| COG4329 | 0.17208 | -0.06077 | 4 | 21 | Membrane |
| 32SB1 | | -0.07736 | 0 | 25 | Ferredoxin |
| COG1020 | 0.09493 | -0.0639 | 9 | 16 | D-alanine [D-alanyl carrier protein] ligase activity |
| COG3569 | 0.08176 | -0.07284 | 3 | 22 | DNA Topoisomerase |
| COG2094 | 0.05304 | -0.10528 | 16 | 9 | Belongs to the DNA glycosylase MPG family |

**Supplementary Table 12**: Feature importance (top 10) for JUICE Korou trait: halophile

HALO

| Feature | Mean SHAP If Present | Mean SHAP If Absent | N(present) | N(absent) | Feature Annotation |
|---|---|---|---|---|---|
| COG0655 | 0 | -0.09704 | 8 | 17 | NAD(P)H dehydrogenase (quinone) activity |
| COG1027 | -0.0604 | 0 | 14 | 11 | Aspartate ammonia-lyase |
| COG1613 | -0.09166 | 0 | 9 | 16 | Sulfate ABC transporter periplasmic sulfate-binding protein |
| COG2733 | -0.10138 | 0 | 8 | 17 | membrane |
| COG1236 | 0 | -0.04036 | 5 | 20 | Exonuclease of the beta-lactamase fold involved in RNA processing |
| COG4967 | 0 | -0.03822 | 5 | 20 | type IV pilus modification protein PilV |
| COG1272 | 0 | -0.06309 | 13 | 12 | protein, Hemolysin III |
| COG1528 | -0.07226 | 0 | 10 | 15 | Iron-storage protein |
| COG0340 | -0.0508 | 0 | 14 | 11 | biotin-[acetyl-CoA-carboxylase] ligase activity |
| COG5006 | 0 | -0.06392 | 14 | 11 | permease, DMT superfamily |

**Supplementary Table 13**: Feature importance (top 10) for JUICE Korou trait: haloresilience

**Haloresilience**

| Feature | Mean SHAP If Present | Mean SHAP If Absent | N(present) | N(absent) | Feature Annotation |
|---|---|---|---|---|---|
| COG1292 | 0.19263 | -0.50909 | 13 | 12 | Belongs to the BCCT transporter (TC 2.A.15) family |
| COG2035 | 0.31954 | -0.227 | 5 | 20 | Membrane |
| COG0471 | 0.15232 | -0.24741 | 10 | 15 | metal ion transport |
| COG3476 | 0.13557 | -0.14149 | 11 | 14 | COG3476 Tryptophan-rich sensory protein (mitochondrial benzodiazepine receptor homolog) |
| COG1320 | 0.07045 | -0.15821 | 9 | 16 | monovalent cation:proton antiporter activity |
| COG0798 | 0.09563 | -0.1274 | 4 | 21 | PFAM Bile acid sodium symporter |
| COG2212 | 0.08372 | -0.12727 | 9 | 16 | antiporter activity |
| COG1863 | 0.06138 | -0.13561 | 11 | 14 | multisubunit Na H antiporter MnhE subunit |
| COG2130 | 0.07798 | -0.08779 | 9 | 16 | NADP-dependent |
| COG0415 | 0.06394 | -0.09823 | 12 | 13 | Belongs to the DNA photolyase family |

**Supplementary Table 14**: Feature importance (top 10) for JUICE Korou trait: psychrophile

**PSYCHRO**

| Feature | Mean SHAP If Present | Mean SHAP If Absent | N(present) | N(absent) | Feature Annotation |
|---|---|---|---|---|---|
| COG0255 | 0 | 0.07767 | 16 | 9 | Belongs to the universal ribosomal protein uL29 family |
| COG0185 | 0 | 0.0794 | 17 | 8 | Protein S19 forms a complex with S13 that binds strongly to the 16S ribosomal |
| COG0096 | 0 | 0.07424 | 17 | 8 | One of the primary rRNA binding proteins, it binds directly to 16S rRNA central |
| COG0736 | 0 | 0.09481 | 19 | 6 | holo-[acyl-carrier-protein] synthase activity |
| COG0097 | 0 | 0.06843 | 17 | 8 | This protein binds to the 23S rRNA, and is important in its secondary structure. |
| COG0344 | 0 | 0.041 | 12 | 13 | acyl-phosphate glycerol-3-phosphate acyltransferase activity |
| COG0416 | 0 | 0.04235 | 13 | 12 | fatty acid biosynthetic process |
| COG2865 | -0.04963 | 0 | 9 | 16 | translation initiation factor activity |
| COG4552 | -0.04527 | 0 | 9 | 16 | Acetyltransferase involved in intracellular survival and related |
| COG0775 | 0 | 0.05595 | 18 | 7 | Catalyzes the irreversible cleavage of the glycosidic bond in both 5'-methylthio |

**Supplementary Table 15**: Feature importance (top 10) for JUICE Korou trait: sporulation

| sporulation | | | | | |
|---|---|---|---|---|---|
| Feature | Mean SHAP If Present | Mean SHAP If Absent | N(present) | N(absent) | Feature Annotation |
| COG3590 | -0.51767 | 0.01956 | 13 | 12 | peptidase |
| COG4521 | 0 | -0.2102 | 8 | 17 | taurine ABC transporter |
| COG4326 | 0.2408 | -0.12369 | 1 | 24 | sporulation control protein |
| COG2971 | 0 | -0.14513 | 6 | 19 | BadF BadG BcrA BcrD |
| COG3448 | 0 | -0.15979 | 10 | 15 | diguanylate cyclase activity |
| COG1988 | 0 | -0.11915 | 5 | 20 | membrane-bound metal-dependent |
| COG0729 | -0.40303 | 0.0126 | 5 | 20 | surface antigen |
| COG0432 | 0 | -0.08221 | 4 | 21 | Pfam Uncharacterised protein family UPF0047 |
| COG0402 | 0 | -0.10443 | 9 | 16 | S-adenosylhomocysteine deaminase activity |
| COG2169 | 0 | -0.0922 | 9 | 16 | Transcriptional regulator |

### **Supplementary Table 16**: Description and performance of PhenDB models.

| Model name | Description | Maximal accuracy |
|---|---|---|
| but | Organism is producing butyric_acid | 0.93 |
| ARCHAEA | Organism is an archaeon (C) | 1 |
| SYMBIONT | Organism has an obligate intracellular lifestyle (C) | 0.99 |
| for | Organism is producing formic_acid | 0.78 |
| sporulation | Organism is capable of producing endospores for persistence (T) | 0.96 |
| AOB | Organism is part of the clade of ammonia-oxidizing bacteria (E) | 0.97 |
| AUTO | Organism is capable of growth with CO2 as sole carbon source (C) | 0.87 |
| lac_L | Organism is producing L_lactic_acid | 0.81 |
| gram_stain | Cells of organism stain Gram-positive (+) or Gram-negative (-) (T) | 0.98 |
| nif | Organism is capable of fixing N2 (E) | 0.92 |
| succ | Organism is producing succinic_acid | 0.9 |
| NOB | Organism is part of the clade of nitrite-oxidizing bacteria (E) | 0.81 |
| glc_D | Organism is utilizing D_glucose | 0.82 |
| h2 | Organism is producing hydrogen | 0.83 |
| T4SS | Organism expresses a Type IV secretion system (C) | 0.91 |
| ac | Organism is producing acetic_acid | 0.77 |
| lac_D | Organism is producing D_lactic_acid | 0.86 |
| phototrophy | Organism captures light energy and convert it into chemical energy | 0.91 |
| indole | Organism is producing indole | 0.83 |
| actn_R | Organism is producing R_acetoin | 0.9 |
| Sulfate_reducer | Organism is capable of using sulfate as terminal electron acceptor (T) | 0.96 |
| THERM | Organism has a thermophilic lifestyle (C) | 0.79 |
| Saccharolytic | Organism has a Saccharolytic lifestyle | 0.91 |
| Aerobe | Organism is capable of aerobic respiration (T) | 0.95 |
| Asaccharolytic | Organism has a Asaccharolytic lifestyle | 0.75 |
| Fermentative | Organism has a Fermentative lifestyle | 0.82 |
| isobut | Organism is producing isobutyric_acid | 0.85 |
| HALO | Organism has a halophilic lifestyle (C) | 0.7 |
| nonfermentative | Organism has a nonfermentative lifestyle | 0.75 |
| T6SS | Organism expresses a Type VI secretion system (C) | 0.95 |
| isoval | Organism is producing isovaleric_acid | 0.91 |
| THERM_OR_TOLERANT | Organism has a thermophilic lifestyle or is thermotolerant (C) | 0.81 |
| btoh | Organism is producing 1_butanol | 0.9 |
| PSYCHRO | Organism has a psychrophilic lifestyle (C) | 0.63 |
| etoh | Organism is producing ethanol | 0.78 |
| METHANOTROPH | Organism is capable of growth with methane as the sole carbon source ( | 0.97 |
| motility | Organism is capable of self-propelled motion (T) | 0.95 |
| Anaerobe | Organism is capable of anaerobic respiration (T) | 0.93 |
| T3SS | Organism expresses a Type III secretion system (C) | 0.93 |
